# Fluid extraction from the left-right organizer uncovers mechanical properties needed for symmetry breaking

**DOI:** 10.1101/2022.04.21.489023

**Authors:** Pedro Sampaio, Sara Pestana, Adán Guerrero, Ivo A. Telley, David. J. Smith, Susana S. Lopes

## Abstract

Humans and other vertebrates define body axis left-right asymmetry in the early stages of embryo development (Shiratori and Hamada, 2006). The mechanism behind left-right establishment is not fully understood (Mizuno *et al*., 2020; Maerker *et al*., 2021; Minegishi *et al*., 2021). Symmetry breaking occurs in a dedicated organ called the left-right organizer (LRO) and involves motile cilia generating fluid-flow therein. However, it has been a matter of debate whether the process of symmetry breaking relies on a chemosensory or a mechanosensory mechanism (Shinohara *et al*., 2012). Novel tailored manipulations for LRO fluid extraction in living zebrafish embryos allowed us to pinpoint a decisive developmental period for breaking left-right symmetry during development. The shortest critical time-window was narrowed to one hour and characterized by a mild counterclockwise flow. The experimental challenge consisted in emptying the LRO of its fluid, abrogating simultaneously flow force and chemical determinants. Our findings revealed an unprecedented recovery capacity of the embryo to re-fil and re-circulate new LRO fluid. The embryos that later developed laterality problems were found to be those that had lower anterior angular velocity and thus less anterior-posterior heterogeneity. Aiming to test if this led to differential endocytosis of any molecular determinant, we replaced the LRO fluid by a physiological buffer. Despite some transitory flow homogenization, laterality defects were absent unless viscosity was altered demonstrating that symmetry breaking does not depend on the nature of the fluid content but rather relies on fluid mechanics. Altogether, we conclude that mechanosensing is the likely mechanism to govern left-right early establishment.

---

Vertebrate organisms break lateral symmetry during embryonic development and define left and right. This left-right axis is the third and final body axis to be established in the embryo and encompasses a biophysical process yet to be fully elucidated. In mouse, frogs, and fish it involves fluid movement in a specialized organ called the left right organizer (LRO). In the mouse embryo, cilia driven fluid flow inside the LRO has been proposed to activate either a mechanosensory mechanism in the epithelial cells that constitute the LRO, or to transport morphogens (as dissolved molecules or carried inside extracellular vesicles) towards the left side of the LRO (Nonaka *et al*., 1998; McGrath *et al*., 2003; Tanaka, Okada and Hirokawa, 2005). While the mechanism is unknown, it culminates in asymmetric left-sided calcium signalling at the node cells and a subsequent conserved Nodal pathway on the left side of the embryo (Tanaka, Okada and Hirokawa, 2005; Takao *et al*., 2013; Mizuno *et al*., 2020). The endpoint of this developmental process is the asymmetric localization of internal visceral organs, such as the heart and liver.

Analysis of cilia driven LRO flow is therefore crucial to understand this initial process of symmetry breaking. The transparent zebrafish embryo is a particularly suitable model system enabling flow visualization and subsequent scoring of organ asymmetry in the same individual. We have shown that wildtype (WT) zebrafish embryos have a stereotyped flow pattern in the LRO (known as Kupffer’s vesicle or KV), exhibiting higher dorsal anterior speed (Sampaio *et al*., 2014; Smith, Montenegro-Johnson and Lopes, 2014). Our group has demonstrated that deviations from this flow pattern due to less cilia or shorter motile cilia, predictably generate larvae with wrong laterality outcomes, indicating a biological relevance to flow (Sampaio *et al*., 2014). Thus, flow dynamics and the mechanical-chemical readout are central to solving the problem of how left-right asymmetry is first established.

Particle tracking for fluid flow reconstruction and modelling have provided quantitative insight in zebrafish (Sampaio *et al*., 2014). Although the LRO is an organ with a limited lifetime, from ~2 ss to 14 ss, seminal work unveiled that the zebrafish Kupffer’s vesicle is dispensable for left-right establishment after 10 somite stages (ss) (Essner *et al*., 2005). Guided by this finding, most subsequent studies have been performed at 8-10 ss, when an asymmetry in *dand5* gene expression is first detected (Lopes *et al*., 2010; Sampaio *et al*., 2014; Ferreira *et al*., 2017; R. Ferreira *et al*., 2019). The clear degradation of *dand5* gene expression on the left side of the LRO is the first asymmetric molecular signal in left-right development, downstream of fluid flow, an event conserved across species (Marques *et al*., 2004; Hojo *et al*., 2007; Schweickert *et al*., 2010).

Yuan and colleagues have shown that, as soon as zebrafish LRO cilia are formed and some start beating, at around 1 to 4 ss, intraciliary calcium oscillations (ICOs) can be registered in LRO cells (Yuan *et al*., 2015). These signals were reported to precede cytosolic calcium elevations that extend mostly to the left mesendodermal regions and were shown to be essential to trigger correct left-right patterning. Therefore, it is conceivable that an asymmetry in *dand5* mRNA expression, first detectable at 8 ss in zebrafish (Lopes *et al*., 2010), results from molecular events that significantly precede that time point. Although it is evident from pioneer studies using mouse mutants without cilia or with immotile cilia that fluid flow has a critical role in conveying a LR biased signal to the nearby cells (Nonaka *et al*., 1998; Supp *et al*., 1999), both the nature of the signal and its detection mechanism remain elusive. The mechanosensory hypothesis postulates that fluid flow can be mechanically sensed by cilia, through a Polycystin protein complex involving PKD1L1 and PKD2 cation channel (Field *et al*., 2011; Kamura *et al*., 2011; Yuan *et al*., 2015). Conversely, the chemosensory hypothesis posits that vesicular parcels or extracellular vesicles containing an unknown chemical determinant are transported by the directional fluid flow towards the left side of the mouse node (Cartwright, Piro and Tuval, 2020). However, the existence of such morphogen has not yet been confirmed or any evidence for its endocytosis has ever been revealed.

To address these 20 year-long-standing questions, we decided to take a step back and unequivocally determine the developmental period for left-right initiation. We envisaged that by determining the critical time-window within the 7 hours lifetime of the zebrafish LRO, we could then characterize its fluid flow and, thus, narrow down the variables that contribute to left-right establishment. Ultimately, we aimed at obtaining evidence for one or the other model for LR initiation as current scenarios fail to fully uncouple the LRO hydrodynamics from the potential role of fluid flow in transporting vesicles or molecules with signalling properties (Cartwright, Piro and Tuval, 2020).

The LRO is filled with fluid, but the nature of the fluid content is unknown because, so far, its minute volume, around one hundred picolitres (Roxo-Rosa *et al*., 2015), prevented a detailed biochemical analysis. However, the swelling mechanism of the LRO lumen is known to be driven by the chloride channel known as cystic fibrosis transmembrane conductance regulator (CFTR) that, when impaired, predictably leads to empty LROs and left-right defects (Navis, Marjoram and Bagnat, 2013). Thus, it is possible to block the swelling process by using pharmacology, anti-sense technology or mutants for CFTR, without affecting cilia number or motility. However, manipulation with precision and higher resolution in time was lacking. To determine the sensitive time-window for LR initiation, we devised a novel assay that allows for highly controlled mechanical manipulation of the LRO fluid. We performed total fluid extractions from the LRO at different developmental time points only 30 minutes apart. After confirming that the KV was still present and able to refill, we followed the development of each embryo until organ laterality could be determined. This approach led us to narrow the time-window for symmetry breaking to one-hour, from 4 to 6 ss, corresponding to the cell rearrangement period first reported by Wang et al. (Wang *et al*., 2011). During this time interval we found fluid flow to be mild but already directional. Extraction of LRO fluid at 5 ss led to 35% of WT embryos with both heart and liver incorrect localization. Embryos did not develop LR defects when the original KV fluid was replaced with Danieau’s buffer, but LR axis was perturbed when viscosity of the same buffer was increased. Altogether, our experiments suggest that fluid flow mechanics, contrary to the nature of fluid content, has a major role in LR initial establishment.

## Results

### Fluid extraction affects LR during one-hour time-window

KV fluid flow was disrupted by performing fluid extraction at each LRO developmental stage using a microscope-based micromanipulation setup connected to microfluidic actuators (Figure 1A). We envisaged that if we found the sensitive time-period, we could then investigate with more precision the properties that were necessary and sufficient to break LR. For this purpose, we challenged WT embryos by extracting the liquid from their LRO lumen from 3 to 12 ss (see extraction Video S1 at 8 ss and Video S2 at 5 ss). Manipulated embryos that recovered the LRO liquid, were later characterized according to heart and liver laterality. This procedure depleted the LRO fluid transiently, disrupting the flow generated by motile cilia, while at the same time it potentially extracted any molecular signals that are present in the fluid.

**Figure 1.**
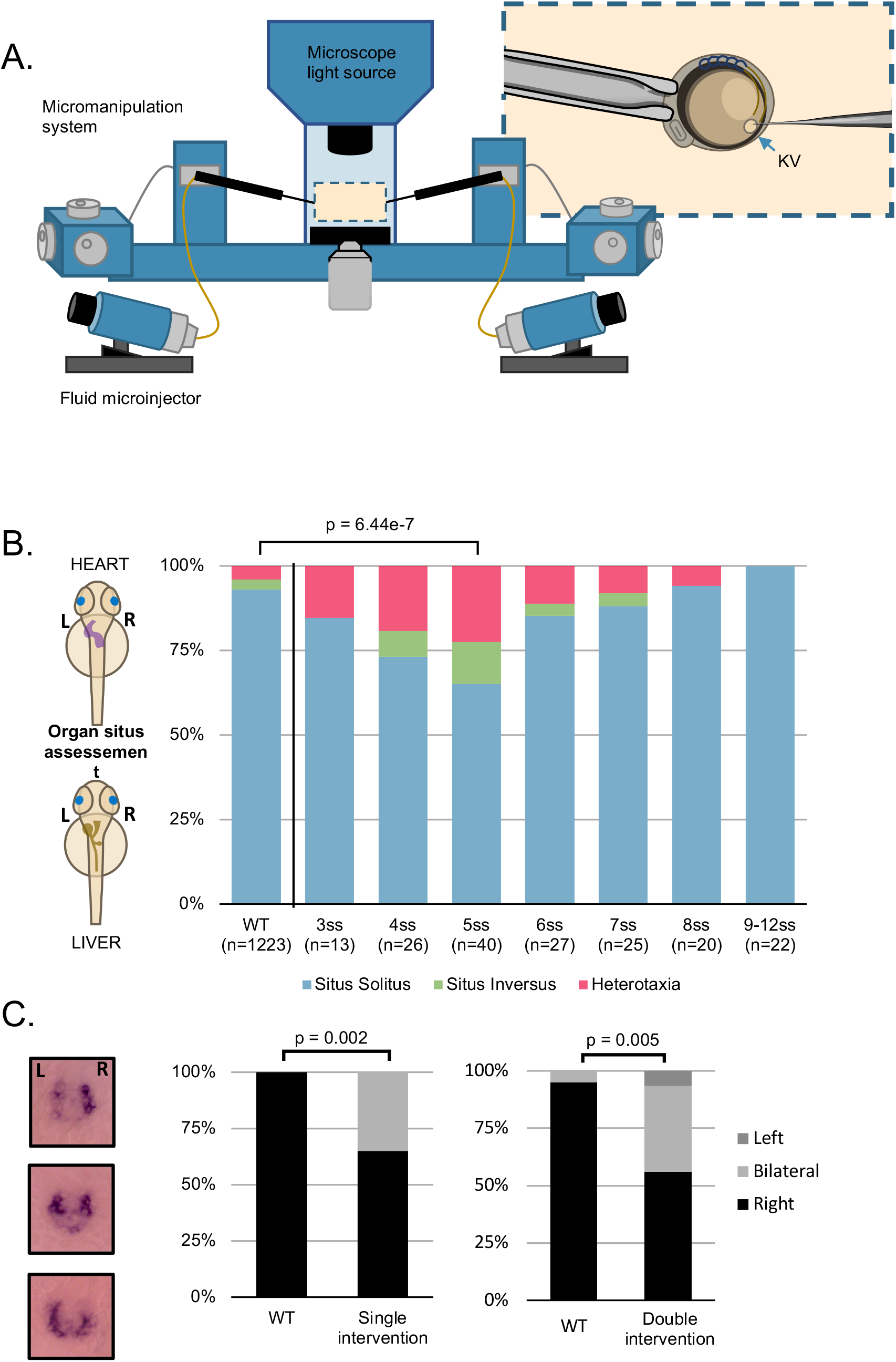
Fluid extractions uncover a sensitive one-hour interval. (A) Schematic representation on the micromanipulation setup developed for the zebrafish LRO fluid extraction throughout development. (B) Evaluation of organ patterning after 48 hours. Heart and liver position were scored for each embryo manipulated in the respective developing stage from 3 to 12 ss, *situs inversus* in zebrafish means both organs are localized on the right side and heterotaxia means that at least one organ is wrongly localized. (C) *dand5* expression pattern at 8 ss after single and double intervention for LRO liquid extraction. LRO. left-right organizer, ss. somite stage.

Controls for these experiments included confirmation that cilia number, length and anterior-posterior (A/P) distribution were not perturbed after fluid extraction at 5 ss, in fixed samples at 8 ss (Figure S1) and that number of motile / immotile cilia and their A/P distribution was not changed when evaluated by live imaging at 6 ss (Figure S2A-D and Table S1). Concomitantly, we analysed cilia beat frequency (CBF) of motile cilia, by high-speed video microscopy. Sham controls, where the LRO was perforated with the extraction needle without applying suction, were included. At 6 ss, no significant difference was found between Sham controls with an average of 38.93 Hz (n = 56 cilia; 6 embryos) and fluid extracted embryos with 37.85 Hz (n = 60 cilia; 6 embryos); (Figure S2B).

To determine if after extraction at 5 ss there were significant anterior-posterior cilia distribution differences between ‘Sham’ and manipulated experimental groups the spatial distribution of cilium types was compared by live imaging as well as the ratio of motile to immotile cilia (Video S3). Consistent with previous studies (Tavares *et al*., 2017), cilia from control ‘Sham’ embryos at 6 ss (n = 7) were predominantly motile (82 ± 4 SEM %) and localized preferentially to the anterior half of the LRO (Fisher test, p-value < 0.05, Figure S2C). The total number of motile cilia for each analysed ‘Sham’ embryo revealed some intrinsic embryo variability even among the control group as reported before (Sampaio *et al*., 2014). Similarly, in the manipulated group (n = 6 embryos), motile cilia were more abundant (78 ± 10 %) and were distributed more anteriorly (Fisher test p-value < 0.05; Figure S2C). Therefore, no significant differences were found for the ratio of motile / immotile cilia between manipulated group compared to ‘Sham’ controls (Figure S2D).

Although most embryos recovered the fluid extraction procedure without LR defects (Figure 1B), some presented both heart and liver misplacements, starting with intervention at 3 ss (2/13 embryos, 15%, p-value = 0.235), becoming more pronounced for intervention at 4 ss (7/26 embryos, 27%, Fisher test, p-value = 0.002) and peaking with intervention at 5 ss (14/40 embryos, 35%; Fisher test, p-value = 6.44e-7). LR organ misplacements then became progressively non-significant from 6 ss onwards (4/27 embryos, 19%; Fisher test, p-value = 0.865; Figure 1B). At this point, as a response to the fluid extraction challenge, we had identified the most sensitive time-window for LR establishment. Next, we wanted to investigate in detail what were the factors that did not allow some of the challenged embryos to recover the correct laterality.

To elucidate how fluid depletion affected LR signalling we examined the expression pattern of *dand5*. It is well established that in WT embryos, zebrafish *dand5* is mainly expressed on the right side of the LRO at 8 ss (Lopes *et al*., 2010) due to left-sided degradation of *dand5* mRNA. Thus, we first evaluated *dand5* expression pattern at 8 ss in embryos that were manipulated at 5 ss, which was the developmental stage with highest incidence of LR defects upon liquid extraction. We observed that *dand5* expression also became abnormally bilateral in 35% of the cases (6/17 manipulated embryos compared to 0/26 controls, Fisher test, p-value = 0.002; Figure 1C) suggesting that either lack of fluid flow or depletion of a molecular signal prevented left-sided degradation of *dand5* mRNA. Next, to further challenge the system we performed an additional fluid extraction one hour after the first intervention when enough liquid was replenished, at 7 ss. Results showed a similar proportion of bilateral *dand5* expression (6/16, 37.5%) to that observed at 5 ss (comparing both plots in Figure 1D and 1E; Fisher test p-value = 0.728). This experiment confirmed that fluid extractions at 7 ss no longer affect LR development significantly, as already implied in Figure 1B. To continue the investigation on a potential deficiency in flow recovery in the ‘LR Defects’ group after fluid extraction at 5 ss, we next assessed how fluid flow dynamics varied with time by retrospectively comparing embryos with and without LR defects.

### LRO recovers lumen area after fluid extraction

The lumen of manipulated LROs started to expand soon after fluid extraction (Figure S3A), indicating that the fluid secretion machinery mediated by CFTR (Navis, Marjoram and Bagnat, 2013; Roxo-Rosa *et al*., 2015) was not collaterally affected by the manipulation procedure and was able to drive the swelling of the LRO lumen. Our observations contrasted with previous reports (Essner *et al*., 2005; Hojo *et al*., 2007) of irreversibly emptied LROs after manipulations, highlighting that the technical approach developed in this work confers minimal damage to the LRO core structure.

Gokey et al.(Gokey *et al*., 2016) showed that below 1300 μm^2^ of KV cross-sectional area at 8 ss, LR patterning could be compromised. Therefore, a scenario where some LROs replenish better than others could explain a differential recovery, enabling the correct organ patterning in most, but not all embryos. Thus, we first investigated if there were significant differences in fluid recovery between manipulated embryos that developed LR defects from those that did not. We performed LRO fluid extraction at 5 ss and followed the recovery of the LRO cross-sectional area and fluid dynamics along development (at 6 ss, 7 ss and 8 ss). As expected, the LROs from the control group (‘Sham’) were larger at each sampled time point (t-test, p-value < 0.05) compared to the fluid extracted groups (Figure S3B). Further, our results showed that the LRO area was similar as well as the area change per somite (t-test, p-value > 0.05) in manipulated embryos that developed a normal organ patterning (‘No Defects’ group) and those that developed incorrect LR axis establishment (‘Defects’ group, (Figure S3B, C).

### Fluid flow angular velocity is reduced in manipulated embryos showing LR developmental defects

Flow can be determined by tracking particles serving as fiduciary markers. The general pattern of fluid flow in the LRO is roughly circular with local variations (Sampaio *et al*., 2014) (see Videos S4 for an embryo that developed normal LR development and Video S5 for an embryo that showed abnormal LR development). Local flow speed (velocity magnitude) is a useful measure for identifying regional flow patterns, but it does not fully describe the directional material transport of the LRO fluid (Ferreira *et al*., 2017; Juan *et al*., 2018). Thus, we determined the angular velocity at discrete locations as a proxy for the circular flow speed (Figure 2A-C). Considering the two sensing hypotheses, we predicted that any type of detectable LR signal, either as flow-induced shear force or as a signalling molecule, must happen in the vicinity of the cell membrane. Taking this into consideration, we analysed the fluid dynamics within the outer half of the radius of the LRO lumen.

**Figure 2.**
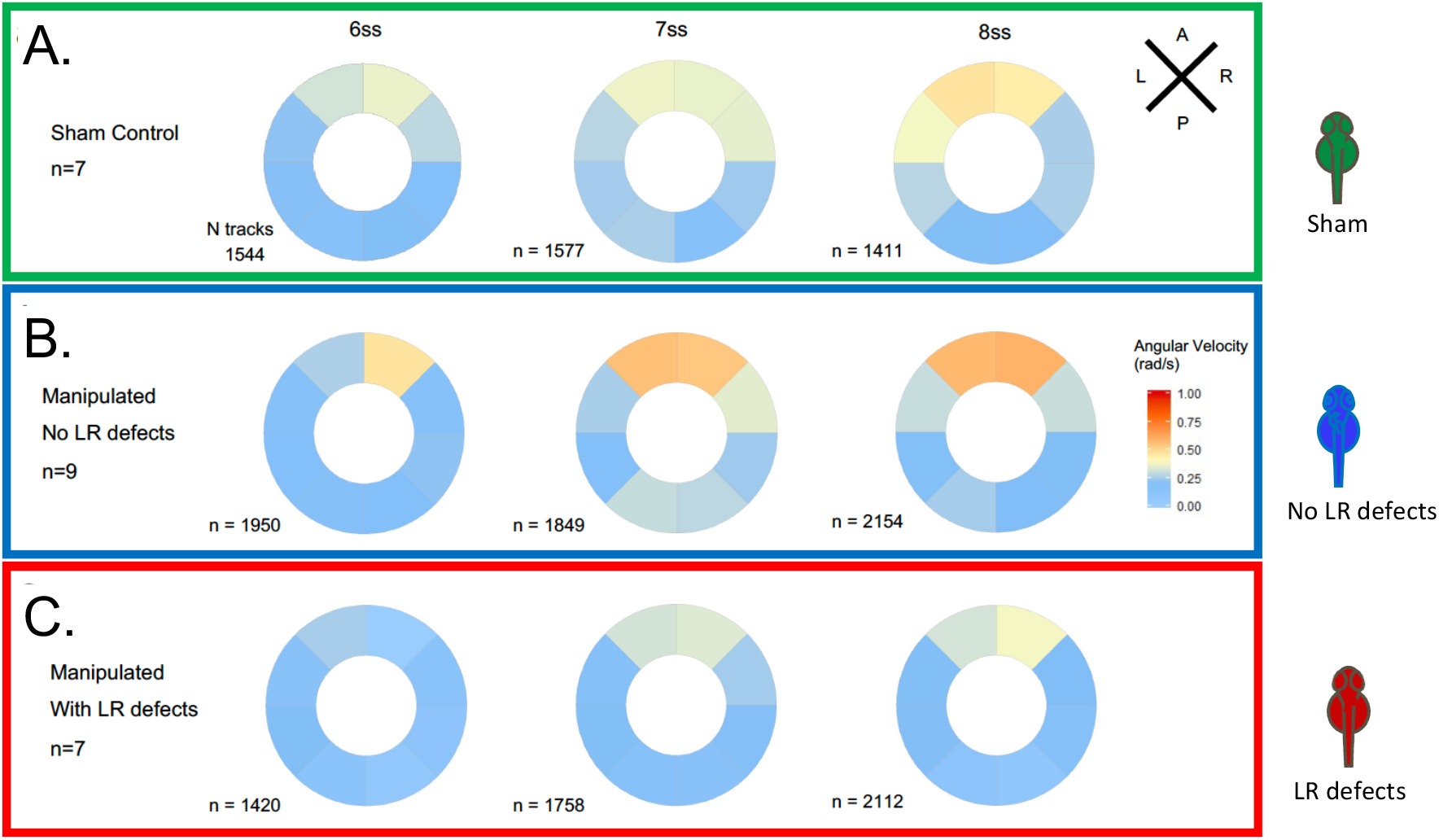
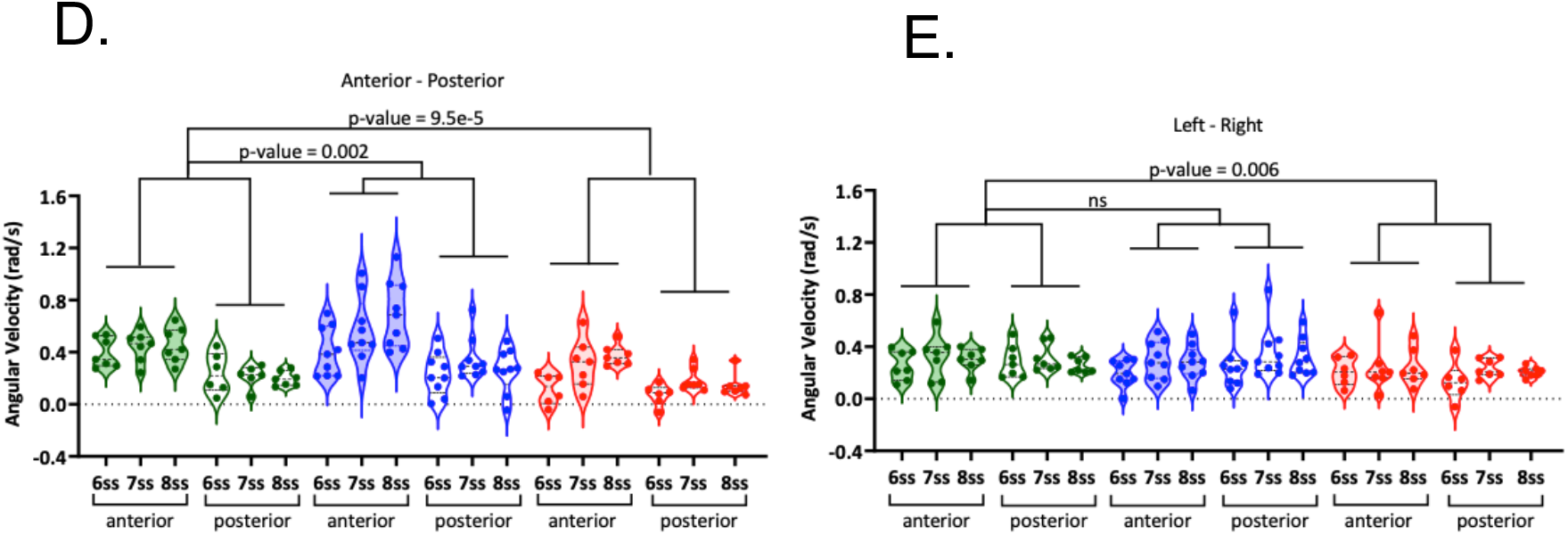
During LRO fluid flow recovery angular velocity decreases in embryos that develop left-right defects. Angular velocity polar plots for 6 ss, 7 ss and 8 ss for the three different groups (A) “Sham” control embryos colour coded in green, (B) “No LR Defects” group of embryos colour coded in blue and (C) “LR Defects” group of embryos colour coded in red. LR defects refer to misplacement of heart or liver encountered at the larval stage after fluid extraction was performed at 5 ss; number of tracks refers to the number of particle trajectories identified for the quantifications and respective angular velocity plots. Colour code on polar plots refers to the median angular velocity for pooled embryos. (D-E) Violin plots show quantifications and statistical comparisons of angular velocities found for the tracks analysed (D) anterior-posterior and (E) left and right. Dots contained in the violin plots correspond to median values per embryo. A statistical linear mixed model was applied.

Angular velocity data from the many tracks obtained from each of the sham and extraction experiments were analysed at 6, 7 and 8 ss, first dividing the LRO in 8 radial sections and plotting the median angular velocity per section (Figure 2A-C). To account for the hierarchical structure of the data with both within- and between-embryo variability in angular velocity tracks, linear mixed effects models were fitted to characterise how angular velocity varied across the LRO and over time (Figure 2D-E), incorporating somite stage, normalized LR/ AP axis position and group (Sham/ Defects/ No Defects) as independent variables. Coefficient values can be interpreted as showing relative effect sizes and direction (see Table 1).

**Table 1.** Linear mixed-effects model fit: Results for fluid extraction experiment. Fixed effects coefficient estimates are given. Model information: Number of observations: 15775; embryos n = 23; Fixed effects coefficients: 12; Random effects coefficients 69; Covariance parameters: 7. Model formula: AAV ~ 1 + Group*LR + Group*PA + Group*Time + (1 + Group | EmbryoID). Results relate to **Figures 2D-E;**\*\*\* indicates p<0.001; ** indicates p<0.01; * indicates p<0.05

| Name | Estimate | Lower CI | Upper CI | p-value |
| --- | --- | --- | --- | --- |
| (Regression line intercept) | 0.2515 | 0.20762 | 0.29544 | 3.8408E-29 *** |
| No defects group | -0.032775 | -0.12321 | 0.05766 | 0.47748 |
| Defects group | -0.12728 | -0.18434 | -0.070215 | 1.2396E-05 *** |
| Left-right axis | 0.0042934 | -0.0092686 | 0.019363 | 0.57655 |
| Posterior-anterior axis | 0.16131 | 0.14636 | 0.17626 | 6.285E-98 *** |
| Somite stage | 0.02145 | 0.012091 | 0.030809 | 7.0847E-06 *** |
| Interaction of no defects group and left-right axis | 0.026653 | 0.007577 | 0.04573 | 0.006176 ** |
| Interaction of defects group and left-right axis | -0.0026553 | -0.022753 | 0.017443 | 0.79567 |
| Interaction of no defects group and posterior-anterior axis | 0.033645 | 0.012118 | 0.055171 | 0.0021912 ** |
| Interaction of defects group and posterior-anterior axis | -0.041406 | -0.06219 | -0.020622 | 9.461E-05 *** |
| Interaction of no defects group and somite stage | 0.026635 | 0.014317 | 0.038953 | 2.2637E-05 *** |
| Interaction of defects group and somite stage | 0.013497 | 0.00056621 | 0.026427 | 0.040778 * |

The linear mixed model for experiments with fluid extraction and ‘Sham’ intervention at 5 ss identified the following significant fixed (embryo-independent) effects: (i) a significant reduction of angular velocity in the ‘Defects group’ (p-value = 1.2e-5, coefficient −0.13 rad/s), (ii) greater angular velocity in the anterior compared with posterior region of the LRO (p-value = 6.3e-98, coefficient 0.16 rad/s), (iii) increased flow velocity over time (p-value = 7.1e-6, coefficient 0.021 rad/s/stage), (iv) enhanced left-right difference in the ‘No Defects group’ (p-value = 0.006, coefficient 0.027 rad/s), (v) reduced posterior-anterior difference in the ‘Defects group’ (p-value = 9.5e-5, coefficient −0.041 rad/s), (vi) enhanced posterior-anterior difference in the ‘No Defects group’ (p-value = 0.002, coefficient 0.034 rad/s), (vii) enhanced degree of velocity increase over time in the ‘No Defects’ group (p-value = 2.3e-5, coefficient 0.027 rad/s/stage) and (viii) (at marginal significance level) some degree of velocity increase over time in the ‘Defects group’ (p-value = 0.041, coefficient 0.013 rad/s/stage).

It is important to note that the increase in angular velocity at the anterior in the ‘No defects’ group compared with the ‘Sham group’ can be explained by the change in size of the Kupffer’s vesicle after manipulation (Figure S3B). Assuming that cilia beating and, therefore, material transport at the periphery is the same in ‘Sham’ group and ‘No-defect’ group (as shown in Figure S2C), the fluid flow absolute velocity is also unchanged, but in a smaller sphere (R = ~25 μm versus R = ~30μm) it leads to a proportionally higher angular velocity. In the ‘Defects’ group, despite this geometric effect, the angular velocity suggests that flow velocity was decreased.

Next, to clarify for further differences between the ‘Defects’ group and the other groups we investigated the directionality of the individual particles as it could provide more cues for localized differences that we observed denoted by negative values for angular velocity indicating the existence of some particles with clockwise flow. We divided the LROs into sections of 30 degrees each and by looking at the direction of each tracked particle in detail (Figures S4-9 and Figure 3A), we only found significant differences in directionality when comparing the group ‘Defects’ with the ‘Sham’ control group in anterior sectors of the LRO at 6 ss (Figure 3B, C). In this region the ‘Defects’ group presented disoriented particle trajectories compared to both the ‘Sham’ control group and the ‘No Defects’ group at 6 ss (Figure S4). This suggests that the manipulation might have regionally perturbed the fluid so that at 6 ss, the earlier time point, the effect of the transient manipulation was still detectable. Conversely, the ‘No Defects’ group never showed significant differences in directionality compared to the ‘Sham’ control group (Figure S4; see also Figure S5 to S9 for complete data on all sectors of the LRO for 7-8 ss).

**Figure 3.**
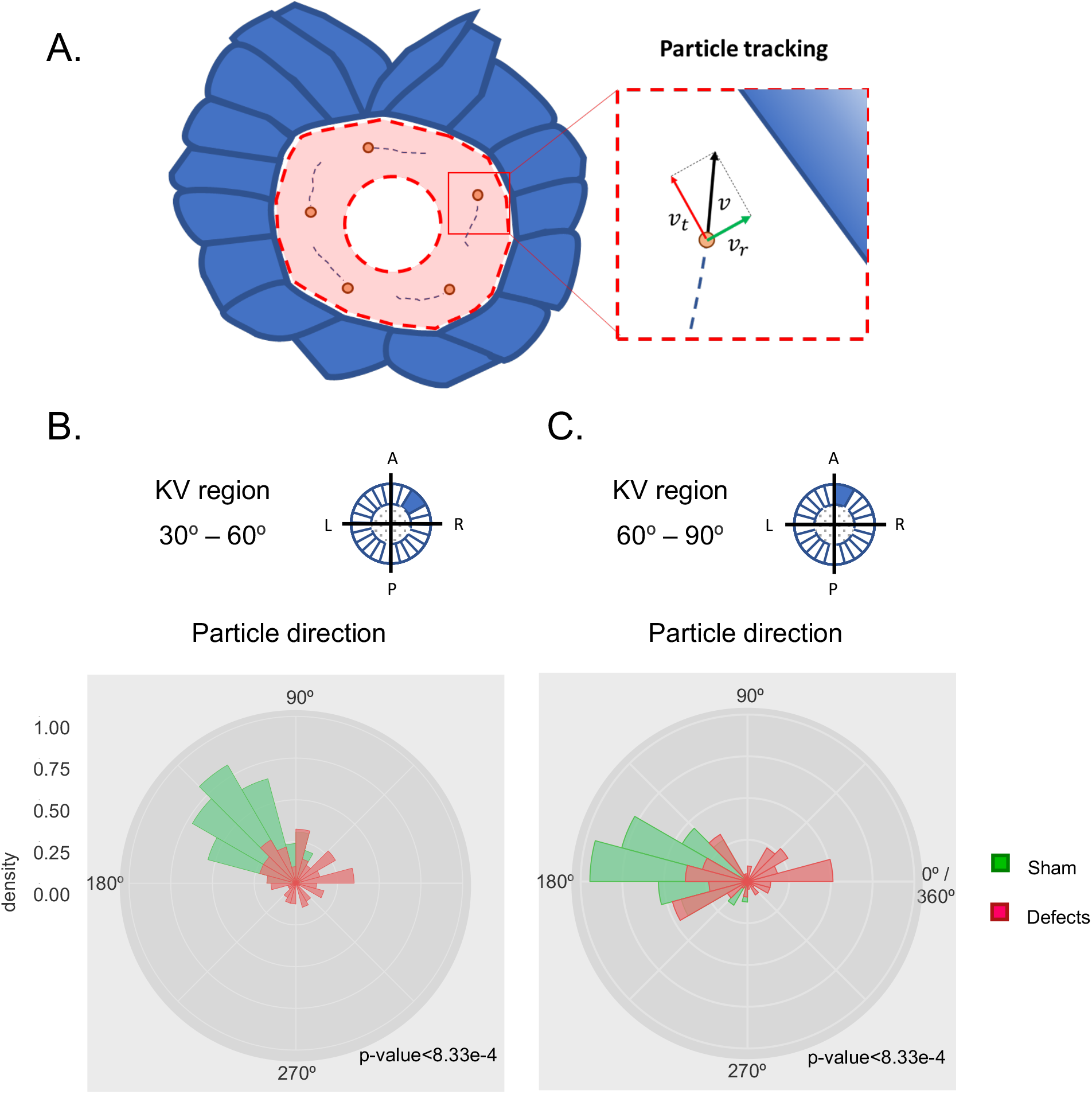
Directionality of vector fields changes in embryos that develop LR defects. (A) Diagram representing particle directionality. LRO area sections were delimited based on intervals of 30 degrees. Highlighted are regions (B) from 30 to 60 degrees and (C) 60 to 90 degrees, that showed significant differences in particle movement between the groups ‘Sham’ control embryos and embryos with ‘LR Defects’. Each density plot represents the pooled tracked trajectory of all moving particles at any given point in time. Respective area region analysed is represented on the top right corner of each plot. To assess differences upon fluid manipulation ‘Sham’ and ‘Defect’ groups were plotted for the 6 ss. Kolmogrov-Smirnov test was used for comparing trajectory distribution between the two groups. Full data can be found in Figures S4-9.

In summary, our data showed that after fluid extraction at 5 ss, the group of embryos that later developed LR defects showed a decreased angular velocity, preferentially at the anterior LRO region resulting in reduced AP difference and it also presented deviations from the counterclockwise flow direction anteriorly. As we could not recognize the region perforated by the needle while analysing angular velocity and particle trajectories, we could not establish causal conclusions regarding needle perturbation in the defects group.

The study of the angular velocity highlighted that a robust flow recovery mechanism is in place in most zebrafish WT embryos, ensuring that LR succeeds in most of the manipulated embryos. Both the groups of embryos that recovered or failed to recover LR development after the extraction manipulation, exposed new sensitive mechanical properties of the LRO flow that will be discussed below.

### A mechanosensory mechanism likely drives LR asymmetry

Extraction of the LRO fluid annihilated both flow dynamics and fluid content, thus it did not allow to discern a potential mechanosensory mechanism from a chemosensory one. Benefitting from the micromanipulation setup we next designed an experiment that could help us to uncouple these two sensory systems. We diluted the fluid content in a physiological buffer (Danieau buffer, DB) without changing the flow dynamics for more than a few seconds, so that we could test whether the fluid content was crucial for LR establishment or not. To address how this dilution experiment affected flow dynamics, we monitored angular velocity and proceeded to its analysis as before from 6 to 8 ss. As a positive control for flow dynamics perturbation, we used methylcellulose (MC, viscosity ~1500 cP; water, viscosity 1 cP) as previously used in *Xenopus* and mouse (Schweickert *et al*., 2007; Shinohara *et al*., 2012) to make the fluid more viscous. We extracted the LRO fluid from 5 ss embryos and diluted it in approximately 1μl volume of Danieau buffer (DB) previously loaded into the needle. Next, after a few seconds, we re-injected the resulting mixed liquid (Figure 4A). To test if dilution of the fluid content was indeed occurring, we performed the same experiment using a rhodamine tracer and confirmed that the LRO lumen became fluorescent after the procedure (Figure 4B). The data for median angular velocities is presented in Figure S10 and the application of a linear mixed model for this set of experiments with DB and MC dilution versus ‘Sham’ intervention at 5 ss identified the following significant fixed effects (Figures 4C-D; Table 2): (i) a significant reduction of angular velocity following methylcellulose (but not DB alone) dilution (p-value = 4.7e-4, coefficient −0.12 rad/s), (ii) greater velocity in the anterior compared to the posterior region of the LRO (p-value = 4.4e-142, coefficient 0.16 rad/s), (iii) increased flow velocity over time (p-value = 5.6e-8, coefficient 0.021 rad/s/stage), (iv) faster flow on the right compared with the left following MC dilution (but not DB alone) (p-value = 0.0039, coefficient 0.026 rad/s), (v) greater difference in flow between anterior and posterior following DB dilution (p-value = 0.0072, coefficient −0.021 rad/s), (vi) less difference in flow between anterior and posterior following MC dilution (p-value = 5.5e-14, coefficient −0.073 rad/s), and (vii) (at the significance level of 0.01) there is some evidence for any positive effect of DB dilution (e.g. in the anterior) receding over time (p-value = 0.010, coefficient −0.013 rad/s/stage). Remarkably, we observed that dilution of LRO fluid with MC significantly reduced cilia beat frequency (31.17 Hz ± 1.4; n = 50 cilia, 5 embryos) as compared to ‘Sham’ controls (39.29 Hz ± 1.2 n = 55 cilia; 5 embryos; Figure 4E, Student’s t-test with Welch’s correction, p-value < 0.01).

**Table 2.**
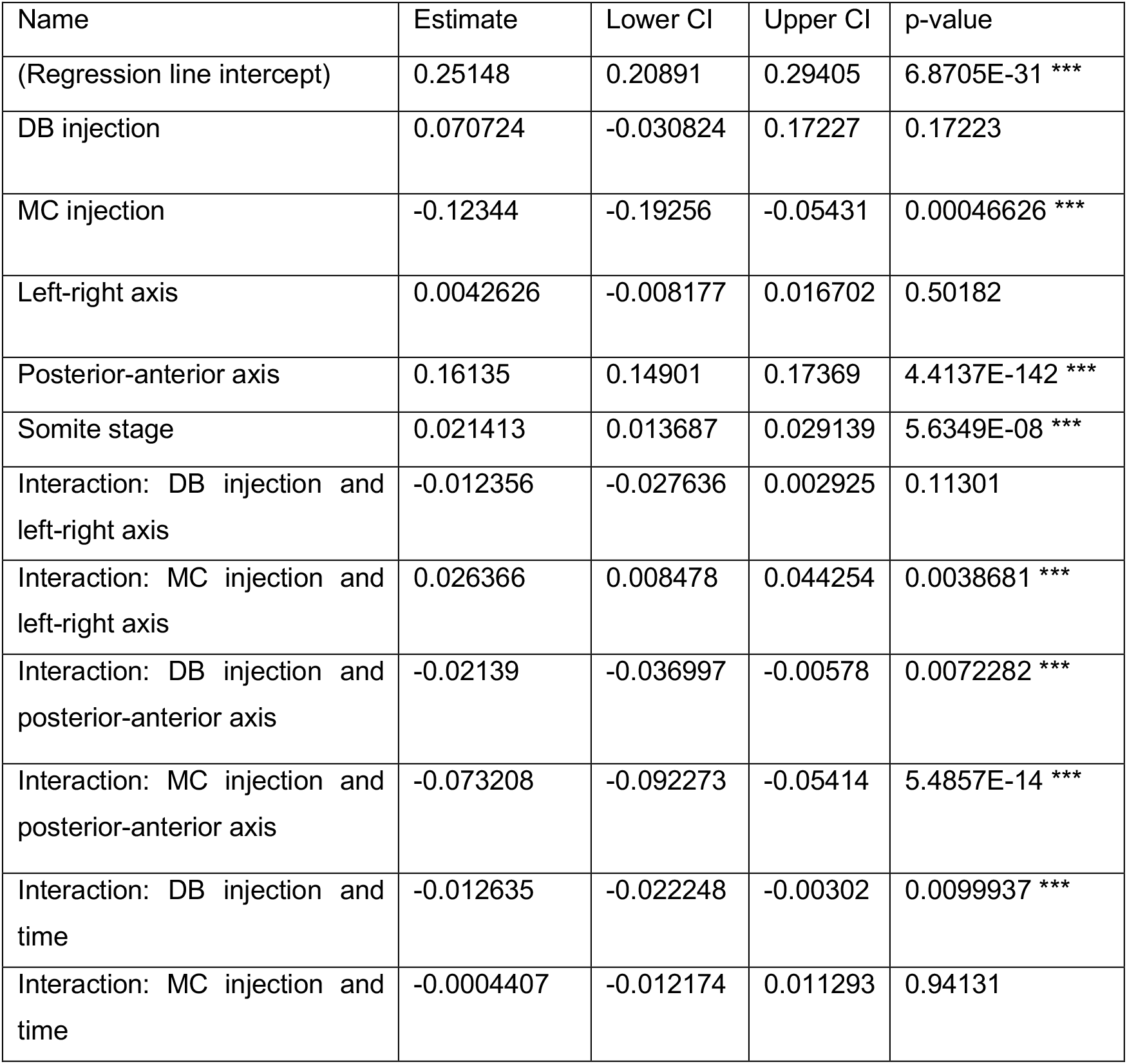
Linear mixed-effects model fit: Results for Danieau’s buffer (DB) and methylcellulose (MC) dilution experiments. Fixed effects coefficient estimates are given. Model information: Number of observations: 16622; n= 30 embryos; Fixed effects coefficients: 12; Random effects coefficients 90; Covariance parameters: 7. Model formula: AAV ~ 1 + Group*LR + Group*PA + Group*Time + (1 + Group | EmbryoID). Results relate to **Figures 4C-D**. Model formula: AAV ~ 1 + Intervention*LR + Intervention*PA + Intervention*Time + (1 + Intervention | EmbryoID). *** indicates p<0.001; ** indicates p<0.01; * indicates p<0.05.

**Figure 4.**
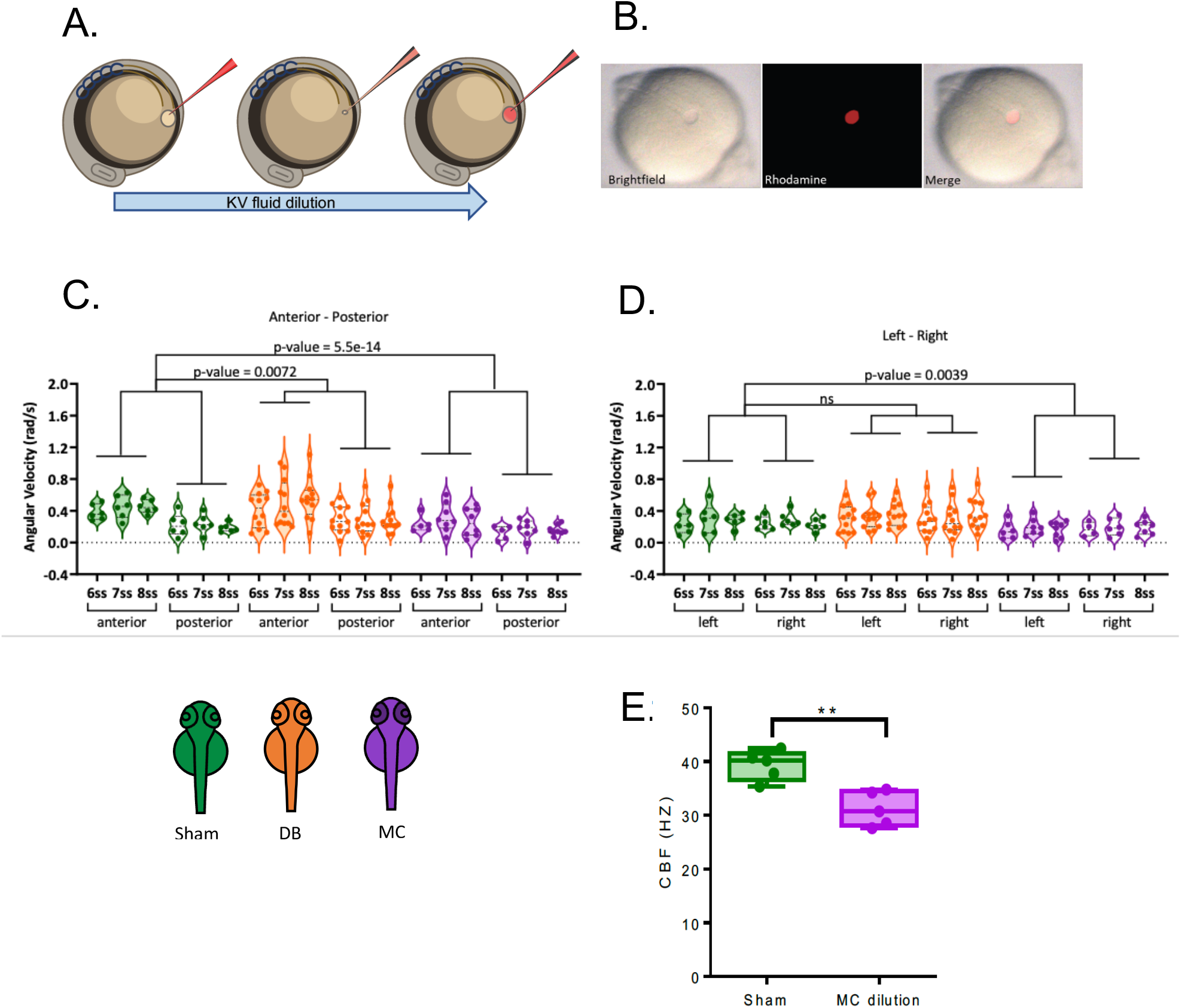
LRO fluid viscosity impacts on angular velocity and cilia beat frequency. (A) Diagram of the dilution experiment: KV fluid is extracted by a needle previously loaded with fluorescent rhodamine-dextran diluted in Danieau’s buffer (DB) that, after mixing of the two liquids, are re-injected into the LRO. (B) Example of a successfully micro-injected embryo labelled with fluorescent rhodamine-dextran. (C-D) Violin plots showing the linear mixed model results for angular velocities (C) anterior-posterior and (D) left and right. Sham control embryos (in green), embryos with LRO fluid diluted with DB (in orange) and embryos diluted with methylcellulose, MC, (in purple). (E) CBF of the motile beating cilia in the ‘Sham’ control embryos versus embryos with LRO fluid diluted with MC. KV: Kupffer’s vesicle; ss: somite stage; DB Danieau’s buffer; MC methylcellulose. A statistical linear mixed model was applied.

Moreover, these experiments demonstrated that following addition of MC but not DB alone, the resulting fluid flow affected *dand5* expression pattern, with more embryos displaying bilateral expression (7/12 embryos, 58%; Figure 5A Fisher test, p-value < 0.001). Consistent results were obtained later in development for organ placement, with 56% of manipulated embryos showing misplacement of the heart position after MC treatment but not DB dilution alone (Figure 5B; Fisher test, p-value < 0.0001).

**Figure 5.**
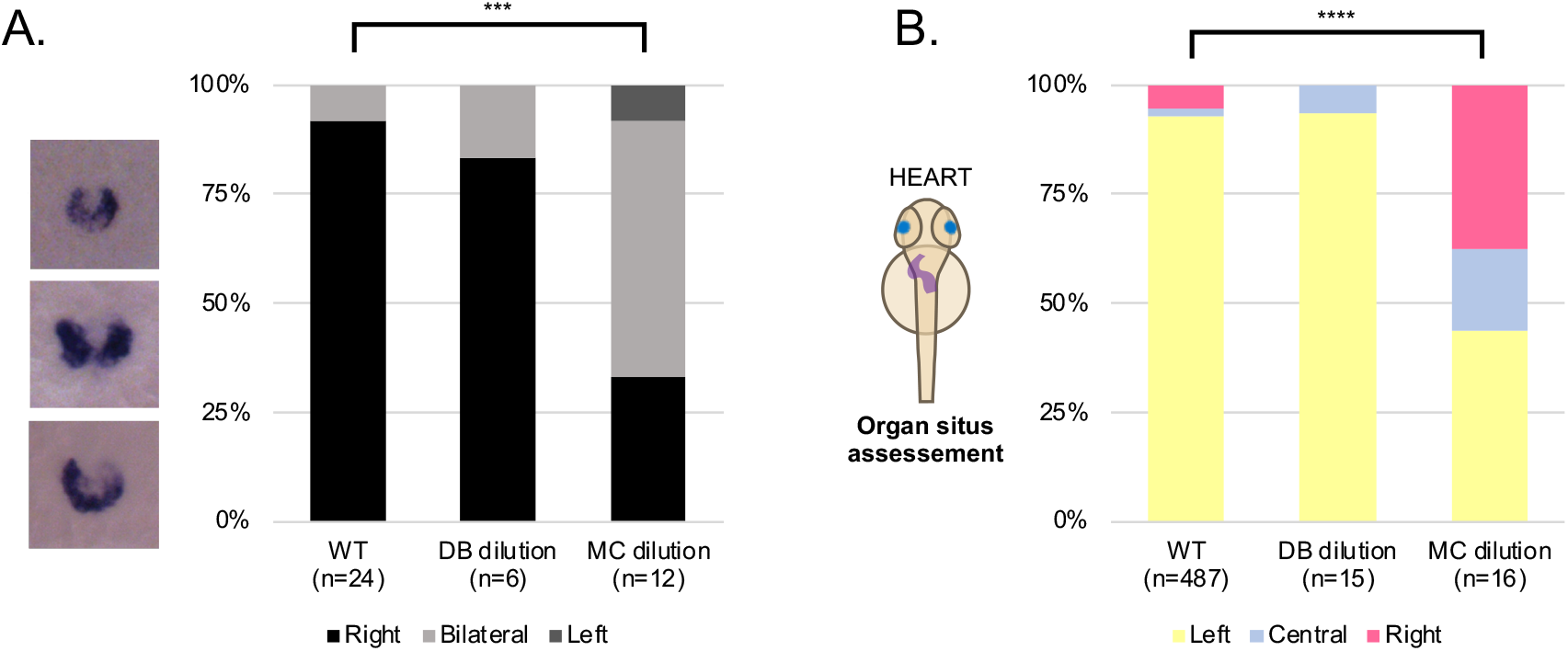
LRO fluid content does not affect left-right development. (A) *dand5* mRNA expression pattern in wild-type control embryos, embryos with LRO fluid diluted with DB and embryos with LRO fluid diluted with MC. Embryos were fixed at 8 ss for *dand5* whole embryo *in situ* hybridization. (B) Heart position (left, central or right sided) was scored at 30 hpf in wild-type control embryos, embryos with LRO fluid diluted with DB and embryos with LRO fluid diluted with MC. ss: somite stage; DB Danieau buffer; MC methylcellulose; hpf: hours post fertilization.

Overall, these experiments provide evidence for flow mechanics in LR development, in support of the mechanosensory mechanism. Indeed, if a signalling molecule contained in the LRO fluid was needed for *dand5* mRNA degradation, then the prediction would be that the DB dilution experiment resulted in many more embryos with *dand5* bilateral expression pattern than it did, as well as more heart misplacements.

Our observation that symmetry breaking is independent of the LRO fluid composition is consistent with the mechanosensory theory. Thus, when liquid volume and fluid dynamics are not changed for more than a few seconds, we showed that LR will be successfully established. Alternatively, if we attempt to interpret these results considering the chemosensory mechanism, dilution of extracellular vesicles (EV) or molecules within could only occur without impacting LR if its secretion rate was sufficiently high to allow a fast turnover, a matter that will be discussed later.

### A mathematical model of flow disruption predicts the experimental data

Our experimental data showed that one hour time-window, from 4 to 6 ss, is the most sensitive period to the fluid extraction challenge, exposing a relevant shorter developmental timing for early LR establishment. In addition, our data also demonstrated that anterior angular velocity is a crucial flow property. To better interpret the intervention results further and elucidate the probable importance of each somite stage in left-right patterning, we constructed a probabilistic mathematical model of how anterior angular velocity, integrated from 3 ss to 8 ss, produces a symmetry-breaking signal. Anterior angular velocity was modelled as a random variable, with mean and variance changing in time, and in response to intervention, based on mixed effects fitting to observed data and previously reported observations of cilia density.

The unknown model parameters were the sensitivity weightings of the LRO to this angular velocity signal at each stage, with normal development occurring provided the integrated weighted signal exceeding a given threshold. Model parameters were then the six weightings relative to this threshold; parameters were fitted to observed results by simulating 1000 virtual LROs for each intervention. Details of the model construction and fitting procedure are given in Supplemental data and Figures S11 and S12.

The best fit for the full model showed good correspondence with observed experimental data (cf. Figure 1B and Figure 6A) in particular, the highest defect rate close to 40% following intervention at 5 ss, and double intervention (5 ss+7 ss) producing a similar defect rate. This model assigned almost all of the stage weighting to 3 ss - 6 ss (Figure 6B). Taking account of the both stage weighting and anterior angular velocity, nearly 90% of the contribution to symmetry breaking occurred in the 4 ss – 6 ss interval (Figure 6C).

**Figure 6.**
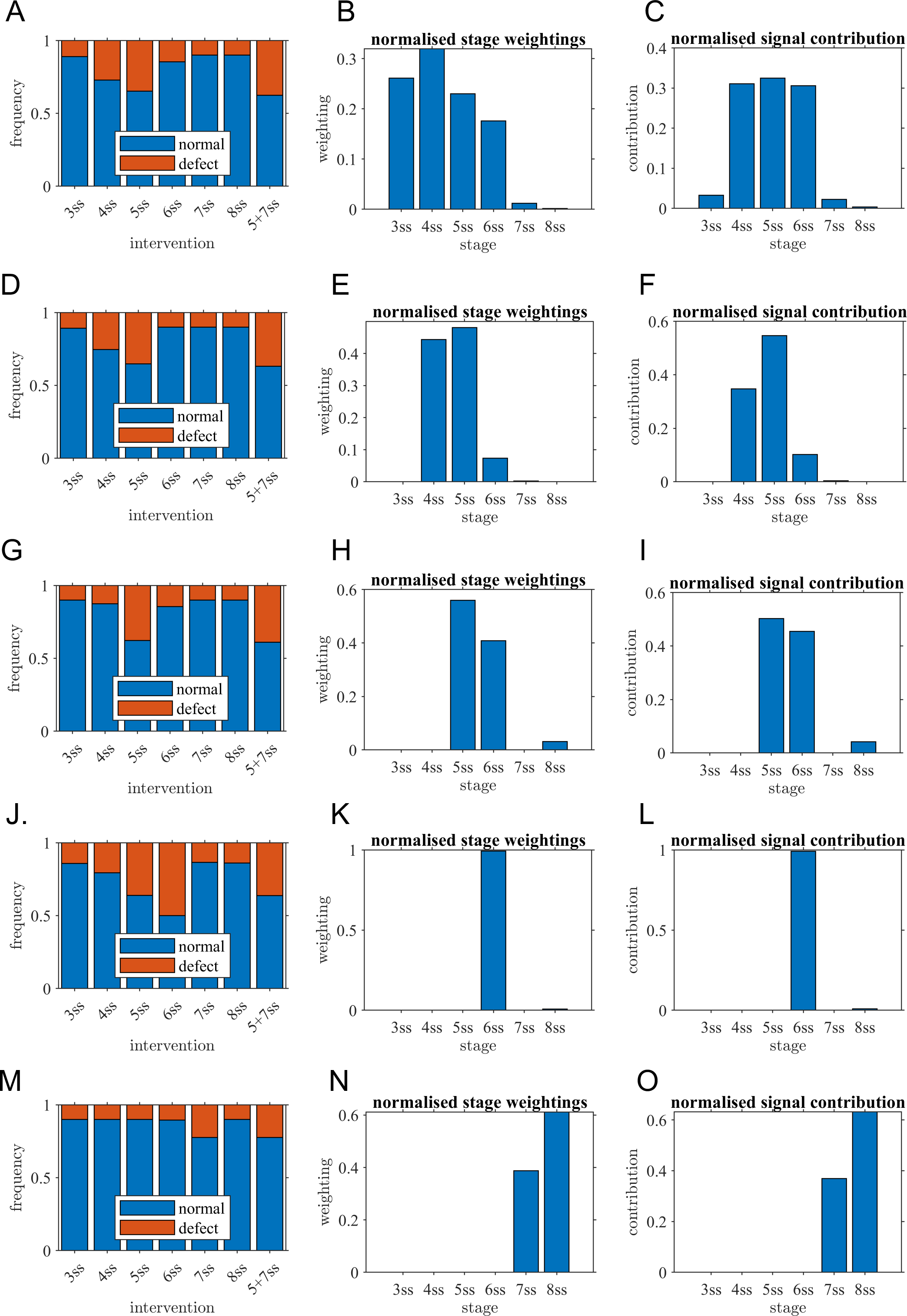
Mathematical modelling confirms the critical role of one-hour interval confirming the experimental observations. (A) full model outputs, fitted to experimental observations in Figure 1B; (B) stage weightings (model parameters *W*(*t*)) for best fit model and (C) stage contributions, calculated by multiplying weightings by average anterior angular velocity. (D-F) model with 3 ss weighting constrained to zero, (G-I) 3 ss - 4 ss weightings constrained to zero, (J-L) 3 ss - 5 ss weightings constrained to zero, (M-O) 3 ss - 6 ss weightings constrained to zero. ss: somite stage.

To assess if models where only later stages were contributing to symmetry breaking could also explain the results, reduced models with each of 3 ss, 3-4 ss, 3-5 ss and 3-6 ss set to zero sensitivity were constructed in a similar manner to assess if observed results could in principle be matched without the contribution of these early stages (Figures 6D-O). The model omitting 3 ss contributions matched observed data similarly to the full model (cf. Figure 1B and Figure 6D). Omitting both 3 ss and 4 ss contributions led to qualitatively similar results (Figure 6G), slightly underpredicting the effect of 4 ss intervention and predicting 5 ss and 5 ss + 7 ss interventions would lead to a higher defect rate than observed. This model assigned almost all the contribution to the 5 ss stage (Figure 6I). Omitting 3 ss - 5 ss contributions led to clearly incorrect predictions of the effect of 6 ss intervention (Figure 6J) and omitting 3 ss - 6 ss contributions led to a model that was not able to make any match to observations (Figure 6M). Overall, these results emphasised the crucial contribution of the 4 ss-6 ss stages, and possible contribution also of the 3 ss stage.

## Discussion

By challenging the zebrafish left-right organizer (LRO) with a new form of manipulation we tested the time limits for the generation of asymmetric signals coming from the LRO, a transient embryonic organ. With this information we established a temporal map for the occurrence of symmetry breaking, highlighting the most sensitive developmental stages and its flow dynamics. We narrowed the LR time-window to 1 hour. After manipulation at 5 ss, we studied flow recovery dynamics from 6 to 8 ss, carefully analysing progression of midplane area, fluid angular velocity components and flow directionality. Most embryos succeeded to display a correct LR organ *situs* and did so by quickly restoring fluid and flow.

In embryos with LR defects we found a general decreased anterior angular velocity, a smaller difference between anterior-posterior angular velocity and an abnormal flow directionality in anterior sections of the LRO. Therefore, we have identified the most sensitive properties of LRO to be placed anteriorly. Yuan et al. (Yuan *et al*., 2015) showed that the intraciliary calcium signal intensity peaked on the left side of the LRO around 3 ss. This period coincides with the first defects we observed upon depletion of the LRO fluid content. Nevertheless, results and modelling showed that these younger embryos may have greater tolerance to challenges and higher chances of recovering the normal LRO flow parameters and subsequent correct LR pattern than those extracted at 5 ss.

Our results showed the most susceptible timing to the fluid manipulation challenge was 4 to 5 ss. Other studies showed that a process of LRO remodelling occurs between 4 and 6 ss resulting in a biased ciliary distribution to the anterior region (Wang *et al*., 2011). It is also at this stage that we previously observed an increased number of motile cilia at the expense of immotile cilia, in robustness of the fluid flow and in establishment of characteristic patterns of fluid dynamics, such as the higher flow speed in the anterior dorsal pole (Smith, Montenegro-Johnson and Lopes, 2014). It would be tempting to suggest that the process of cell remodelling is a landmark after which the LRO becomes less sensitive to challenges. If the challenge is made until 6 ss we demonstrated here that there is evidence for an active sensory mechanism that restores fluid and flow. This heightens the need of identifying the mechanism that senses fluid flow magnitude. Such sensory mechanism seems to be sensitive to flow properties, as viscosity. In the presence of a low Reynolds number, when viscosity dominates and inertia is negligible, a sensory system to be efficient must be very sensitive to lower flow magnitude as suggested in (Shinohara *et al*., 2012). The most plausible mechanosensory system in the zebrafish LRO involves Pkd1l1 and Pkd2, both present in the LRO cilia as shown by immunofluorescence (Roxo-Rosa and Santos Lopes, 2020; Jacinto *et al*., 2021). Although it has not been possible to demonstrate that these two polycystin proteins function together as a mechanosensory complex in zebrafish, there is evidence to be the case in the mouse node (Field *et al*., 2011; Mizuno *et al*., 2020). Moreover, patients with mutations in PKD1L1 or in PKD2 present laterality phenotypes such as *situs inversus*, heterotaxia and congenital heart disease (Bataille *et al*., 2011; Le Fevre *et al*., 2020).

Live quantifications of cilia motility along time, using 2-photon microscopy in Tavares et al. (Tavares *et al*., 2017) attested that in the first stages after the formation of the LRO, there are many more immotile cilia (around 94%) than motile, but then, a gradual loss of cilia immotility occurs during development (Tavares *et al*., 2017). The present study showed that in all embryos that developed correct LR (‘Sham’ and ‘No defects’ groups; scored at 6 ss, very few immotile cilia were present (9 on average) in a total average of 43 cilia (n = 13 embryos), which is only 20,9% of the total LRO cilia. So, according to Ferreira et al. (Ferreira *et al*., 2017), this fact brings to light that perhaps immotile cilia number is not a crucial factor after all. However if we focus the analysis on motile cilia number, our current findings are compatible with a previous fluid mechanics model for the LRO (Sampaio *et al*., 2014) where we estimated that, at least 30 motile cilia are needed for a correct LR outcome.

A more recent hypothesis for the sensory system in the zebrafish embryo postulates that cilia themselves may sense their own movement (Ferreira *et al*., 2019). In fact, Yuan et al. (Yuan *et al*., 2015) filmed intraciliary calcium oscillations (ICOs) in the zebrafish LRO motile cilia in action and showed that ICOs were abrogated in knockdowns for Pkd2 or for cilia motility genes. The motile cilia self-sensing model becomes appealing in view of the flow response to increased viscosity. Motile cilia pattern and power are known to differ at increasing viscosities (Kikuchi *et al*., 2004; Wilson *et al*., 2015; Gallagher *et al*., 2019), which could potentially be detected by a ciliary mechanosensitive transduction channel, like in the fish lateral line where stereocilia bending allows sensing the velocity and direction of water flow (Asadnia *et al*., 2016). In summary, we suggest that a ciliary mechanosensory system, such as Pkd1l1/ Pkd2 could be sensitive to flow magnitude. The fact that CBF is significantly lower in the methylcellulose injected embryos suggests that may be there is a minimum CBF threshold for the sensory system to generate efficient ICOs. Then, how such ICOs could lead to *dand5* mRNA degradation specifically on the left cells of the LRO is not resolved in zebrafish, but according to Yuan et al. (Yuan *et al*., 2015) there is an increase of left-sided ICOs in the early time points that seems to coincide with the crucial time-window we have now established. However, our data on angular velocity does not show a consistent difference in left versus right angular velocity and correct LR development, leading us to suggest that anterior flow is the relevant factor.

Research on LR establishment is moving forward regarding the role of calcium using the mouse model. A previous study by Delling et al.(Delling *et al*., 2016) contradicted the idea that crown cells LRO cilia are good calcium-responsive mechanosensors. Conversely, recent work from Hamada’s lab (Minegishi *et al*., 2021) demonstrated that LR asymmetric intraciliary calcium transients can be in fact detected at the node. Mizuno and colleagues further revealed that discrepancies between the two studies relate to the culture conditions used, which led calcium signals to be missed in the work by Delling et al.(Delling *et al*., 2016). Moreover, recently, Minegishi et al. unveiled that in the mouse model, the 3′-UTR of *dand5* mRNA responds to the direction of fluid flow in a Pkd2-dependent manner via stimulating Bicc1 and Ccr4-Not. Together, these molecular players seem to mediate *dand5* mRNA degradation specifically on the left side of the mouse node and xenopus gastrocel roof plate (Maerker *et al*., 2021). Our independent fluid manipulations support a very sensitive mechanosensory system for the initiation of the LR pathway, but it remains to be demonstrated how and what exactly is sensed by the system. Our data highlights anterior flow magnitude as a major mechanical property in LR. Interventions that decreased flow magnitude or reduced its A/P difference were the ones that generated abnormal LR outcomes.

The time lag and complexity of the recovery process motivated the development of a mathematical model of flow and its developmental influence, and the effect of experimental interventions. Statistical modelling of normal and disrupted flow, and fitting of a ‘weighted contribution’ model to infer the relative importance of each somite stage in explaining the results, confirmed the critical importance of 4 ss to 6 ss in symmetry breaking, and the limited role for flow during 7 ss and 8 ss. The process of constructing the model also emphasised that intervention has two effects: the transient gross reduction in flow associated with removal of the entire luminal fluid, but also a greater variance of flow following recovery – hence observed defects are as much a consequence of a proportion of embryos making a below average recovery, as the interruption itself.

Finally, we need to discuss whether our experiments exclude the potential contribution of molecule or EV secretion to the LR establishment. According to (Ferreira *et al*., 2017) in a simulated transport of signaling molecules in the Kupffer’s vesicle between 9-14 ss for predicted vesicle secretion, left-right concentration gradient and mixing would take 8 seconds to occur, so we predict that earlier at 4-5 ss this process could take longer because there is less flow, perhaps in the order of minutes. Thus, according to this line of thought it is possible that secretion can still occur between our fluid extraction at 5 ss and the limit of the responsive time-window by 7 ss (1 hour interval). Literature on EV secretion rates is scarce but we found an estimate of 60-80 exosomes per cell per hour is expected, depending on the cell type (Agarwal *et al*., 2015; Chiu *et al*., 2016). Our future efforts will be directed to test the role of secretion and endocytosis in the LR symmetry breaking without disrupting fluid mechanics.

## Methods

### Fish husbandry and strains

Zebrafish were maintained at 25°C or 30°C and staged as described elsewhere(Kimmel *et al*., 1995) according to the number of somites. The following zebrafish lines were used for this work: AB and Tg(sox17:GFP)s870(Chung and Stainier, 2008). Procedures with zebrafish were approved by the Portuguese DGAV (Direção Geral de Alimentação e Veterinária).

### LRO manipulation setup for liquid extraction

LRO manipulations of fluid extraction were performed from 3 to 12 somite stages (ss). Zebrafish embryos were individually chosen at the developmental time correspondent to each somite-stage. Furthermore, only embryos that recovered LRO fluid were used throughout the study. Individually embryos were hold from the anterior dorsal side of the body using a holding pipette while a second pipette was used to penetrate the LRO from the posterior dorsal side. After extracting the LRO liquid the microinjection pipette was withdrawn, and embryos were incubated and let to develop. Collectively 173 embryos were manipulated along the different stages. Embryos were let to develop at 28°C and were later characterized according to *dand5* expression pattern or to organ laterality. The micromanipulation system, as illustrated in Figure 1, was composed of an inverted microscope (Andor Revolution WD, Oxford Instruments; or Nikon Eclipse Ti-U) with a 10x Plan DL air objective, a 60x Plan Apo VC water immersion and a 100x Plan Fluor oil immersion objective (Nikon), a set of two motorized axis manipulators (MPC-385-2, Sutter Instruments) placed on both sides of a culture dish for moving pipettes to the desired positions, a CMOS camera (Monochrome UI3370CP-M-GL, Imaging Development Systems GmbH; or high speed FASTCAM MC2 camera, Photron Europe, Limited) to capture images in the microscope field of view, and a set of two CellTram’s (CellTram Vario, FemtoJet, Eppendorf). Each respective microinjector are connected to a holding micropipette for immobilizing the zebrafish embryo, and injection micropipette that allows fluid manipulation from or to the Kupffer’s vesicle (KV). All the setup components were constructed over an optical table with vibration isolation (ThorLabs) to minimize the effects of any residual vibrations or disturbances introduced during the embryo manipulation procedure.

#### Immobilizing embryos

The embryo holding step was based on immobilization of the embryo by a pipette positioned on its left side in order to allow the injection micropipette to be introduced from the right side through the dorsal tissue into the KV. Of the several positions tested immobilization by the zone dorsal to the developing head of the embryo was the most effective. Unlike the yolk, the back of the head offered a resistant tissue with a shape that easily adapted to the diameter of the holding pipette. At the same time this setup allowed the KV to be aligned with the microinjection needle field of action. The creation of a chamber for easy handling of the injection micropipettes while keeping the specimen in an aqueous solution was crucial for the success of this procedure. Using silicone grease (Dow Corning) a rectangular coating was outlined on a coverslip. Next, a sufficient amount of embryo medium E3 was carefully pipetted on the coverslip forming a liquid pond. This setup contained zebrafish embryos in an aqueous environment with very low physical impediment, allowing a high range of contact angle for micropipette manipulation to happen.

#### Pipette forging for micropipette extractions

During its developmental time period the LRO or KV will migrate from 70 to 120 μm below the tail mesodermal cell layer(Kimmel *et al*., 1995). Consequently, vesicular fluid extraction and compound microinjection procedures require resistant sharp capillaries capable of piercing through the tail tissue dorsal to the KV. Micropipette tip size and shape of the taper are crucial factors determining if a micropipette will effectively impale the embryo tissue. Small tips and uniform tapers have the great advantage of causing less cell damage (Sutter Instrument, pipette cookbook, 2008) http://www.sutter.com/contact/faqs/pipette_cookbook.pdf. Likewise, a shorter tapered and very stiff micropipette is needed to penetrate very tough and rigid membranes.

Aluminosilicate glass capillaries are known to have an increased hardness and ability to form small tips compared with its borosilicate counterpart, allowing to form stiff longer tapers important for reaching the KV and avoiding unwanted tissue damage.

Taking this in consideration, microinjection pipettes (Aluminosilicate, inner diameter 0.68mm, outer diameter 1.00, length 10mm, without inner filament) were designed using a custom-made program in a horizontal filament puller (P-97 Micropipette Puller, Sutter Instrument). Reduced cone angle and slender taper can be achieved using a 2-loop puller program.

The resulting pulled pipettes were often sealed or have a very narrow tip (<1 μm). Shaping the tip to the desirable size was important to avoid clogging of the pipette and deliver a sharp cut when impaling the embryo tissue. Therefore, the tips of micropipettes were shaped to small apertures (<5 μm) under the microscope. Using the motorized manipulators, the tips were carefully cut-opened against a polished borosilicate capillary. This allows to have a balance between micropipette longevity and minimal tissue damage during embryo manipulation. For better results reducing the amount of mechanical stress caused by the needle injection, a micropipette beveller (BV-10, Sutter Instrument) was used to increase sharpness of the pipette tips. This additional step was not always carried out due to the significant time increase in preparation of each micropipette.

#### Holding pipette

Since our organism of interest is immersed in an aqueous non-viscous solution, oscillations of the zebrafish embryo are a major concern during micromanipulation. Thus, special care should be taken to assure a rigid stable connection between the embryo tissue and the tip of the micropipette. To achieve this a holding pipette (typically positioned on the left side of the preparation), immobilized the specimen while an opposing injection micropipette (on the right side) was introduced into the embryo tail.

For forging holding pipettes, thin wall borosilicate capillaries (inner diameter 0.75mm, outer diameter 1.00mm, length 10mm, without inner filament) were used. Ideal holding pipettes have a flat straight break at the tip with large inner and outer diameter to provide better attachment and support to the zebrafish embryo. Fire-polishing was then performed to create a smooth surface to interface with the embryo tissue without damaging it, and produce an inner diameter (around 0.50mm) best suited to hold the embryo as described in(Brown and Flaming, 1974), where a common Bunsen burner or a micro forge (MF-900, Narishige) were used to fire-polish the holding pipette.

### Cilia number, length and anterior-posterior distribution in fixed samples

To ensure that low damage is caused during embryo manipulation we further quantified KV cilia number and length upon fluid extraction. After manipulation embryos were immediately fixed in PFA 4% and whole-mount immunostaining for acetylated α-tubulin was performed to label KV cilia as previously described(Lopes *et al*., 2010). Antibodies used for immunostaining were mouse anti-acetylated alpha-tubulin (1:400; Sigma) and and Alexa Fluor 488 (Invitrogen; 1:500). Cilia count and length measurements were performed either manually or through semiautomated detection in IMARIS software program.

### Evaluating motile / immotile cilia distribution by live imaging

Procedure followed the method described in(Tavares *et al*., 2017). Live imaging was performed in a Zeiss LSM 710 confocal microscope with an Olympus 40x water immersion lens (NA 0.8) at room temperature. Cilia identification was performed either manually or through semi-automated detection in IMARIS software program (Bitplane, UK).

### Live imaging of fluid flow in the zebrafish left-right organizer

For the LRO fluid extraction experiments, embryos were maintained between 28°C - 30°C until 5 ss, then mounted in a 2% (w/v) agarose mold and covered with E3 medium, and filmed at 25°C. To evaluate the fluid flow and LRO inflation recovery dynamics, the LRO was followed along development, from 6 to 8 ss in a sample of embryos manipulated at 5 ss (n = 16). In parallel, embryos (n = 6) in which the LRO was punctured by a micropipette, without further liquid extraction, were used as controls (‘Sham’ group). Imaging was performed on a light microscope (Nikon Eclipse Ti-U inverted microscope), under a 60x/1.2 NA water immersion or 100x/1.30 NA oil immersion objective lens, by a FASTCAM-MC2 camera (Photron Europe Limited, UK) controlled with Photron FASTCAM Viewer software. Using the ImageJ plugin Measure Stack, the LRO was delineated, and its luminal area was measured in all focal planes. LRO particles were tracked using the ImageJ plugin MTrackJ as previously reported(Sampaio *et al*., 2014).

### Evaluation of heart and gut laterality

Heart jogging was evaluated at 30 hours post fertilization (hpf) using a stereoscopic zoom microscope (SMZ745, Nikon Corporation). These embryos were then allowed to develop at 28°C and at 53 hpf embryos were fixed and processed for *foxa3 in situ* hybridizations to assess gut laterality(Sampaio *et al*., 2014), or readily scored if a sox17:GFP transgenic line was being used for the respective assay. We could then pair the heart *situs* with gut *situs* for each treatment and attribute an embryo *situs*. Final organ *situs* score was attributed regarding the combined heart and liver lateralization status: *situs solitus* – normal organ patterning (left heart and liver), *situs inversus* - organs positioned opposite to normal positions (right heart and liver), and heterotaxia – comprising all the other laterality defects variants. For the LRO fluid extraction assay a total of 1223 WT embryos and 173 manipulated embryos were characterized according to organ *situs*.

### RNA in situ hybridization and immunofluorescence

Whole-mount *in situ* hybridization and immunostaining was carried out as before(Lopes *et al*., 2010) Digoxigenin RNA probes were synthesized from DNA templates of *dand5*(Hashimoto *et al*., 2004).

### Flow speed, angular velocity and particle directionality

The flow speed, angular velocities and velocity components were calculated by customized scripts in the program environment R (Supplementary data). The centre of the LRO was used as reference point of a polar coordinate system in which angles are expressed in radians. To study the flow dynamics near the LRO apical membrane, tracked points near the centre (corresponding to a circle with half of the LRO radius) were removed, as angular quantities are poorly defined along the axis of rotation, close to the centre of the LRO. Angular velocity represents the effective circular flow magnitude of a particle moving between two consecutive time frames. The centre of the KV was used as reference point of a polar coordinate system in which angles are expressed in radians. Instantaneous angular velocity (rad/sec) was calculated by dividing particle angle change by the time between consecutive image frames (0.2 seconds).

Particle directionality quantification at any given point was obtained by dividing the LRO in 30 degree sections and calculating the change of particle angle between two consecutive time frames, where the axis of the reference is the position of the particle in the first time point. The particle directionality values range from 0 to 2π.

### Fluid dilution and viscosity manipulation in the zebrafish LRO

The KV fluid dilution assay was performed by aspirating the vesicular fluid into a micropipette previously filled with Danieau’s solution 1x (58mM NaCl, 0.7mM KCl, 0.4mM MgSO4, 0.6 mM Ca(NO3)2, 5.0 mM HEPES pH 7.6) diluted in 10,000 MW rhodamine-dextran solution (1:4; Sigma-Aldrich). The mixed solution was injected again into the KV and only embryos with rhodamine-positive KV’s were selected for the experiment. For viscosity manipulation, the micropipette was filled with Danieau’s solution 1x containing 1.5% (w/v) of methylcellulose (M0555, Sigma). Aspirated KV fluid and methylcellulose were then injected again into the KV lumen until reaching the KV volume observed before the manipulation.

### Statistical analysis

A Fisher’s exact test performed with RStudio was used to compare frequency of left-right defects and *dand5* expression patterns between WT and manipulated embryos. To assess potential differences in KV area, cilia number, cilia length between WT and manipulated embryos Student’s t-test was used. To assess area changes per somite Mann-Whitney U test was used. To assess cilia motility differences between anterior and posterior regions Fisher’s exact test was used. Kolmogrov-Smirnov test was used from comparing particle trajectory distribution between two groups. Angular velocity data for tracks imaged in each of the suction and injection experiments were fitted to linear mixed effects models to characterise how angular velocity varies across LRO and over time, how these velocity field patterns are altered due to intervention, and how they are reflected by different embryo fates. Fixed effects were group (injection experiment: sham, buffer injection, methylcellulose; suction experiment: sham, no-defects, defects), somite stage, normalised posterior-anterior axis coordinate, normalised left-right axis coordinate, and interactions of group with somite stage, posterior-anterior and left-right coordinates. Random effects were included to account for heterogeneity between embryos in their response to intervention. The models were specified in Matlab (Mathworks) notation as,

Suction experiment:

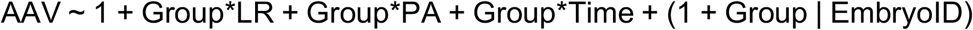

Injection experiment:

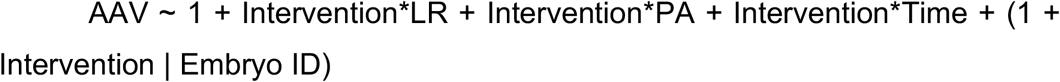

Fitting was carried out with the command fitlme, with effects reported in the main text where p<0.05 and interpreted based on the sign and magnitude of the fitted coefficient.

### Mathematical modelling

Mathematical models of anterior angular velocity, its disruption by suction intervention, and the signal integration process, were implemented as custom scripts in Matlab (R2021a, Mathworks, Inc, Natick MA).

#### Statistical models for anterior angular velocity

The key hypothesis of the model is that symmetry breaking is produced by anterior angular velocity AAV(t) at somite stage *t* integrated from 3 ss to 8 ss, weighted by the a priori unknown sensitivities W(t) at each stage. As the aim here is only to provide a good fit to the flow velocity data as input to the signal transduction model, the precise mathematical form of the model is not relevant. Mixed effects models were fitted to anterior angular velocity data for 5 ss-intervention and sham intervention using the Matlab function nlmefit (nonlinear mixed-effects estimation).

AAV following intervention at 5 ss was modelled using the nonlinear function

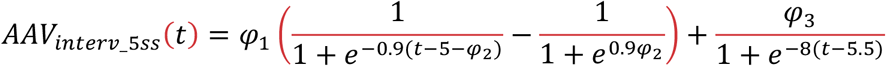

The coefficients 0.9 and 8 were chosen to provide a smooth increase from zero, and the parameters *φ_j_* were determined by fitting. Mixed effects fitting produces estimates for the fixed effect (central tendency in the population) and the standard deviation of the random effect (quantifying variability in the population; to maintain parsimony only the final parameter *φ*_3_ was modelled with a random effect. Estimates for the fixed effects were *φ*_1_ = 0.778 ± 0.679, *φ*_2_ = 0.969 ± 2.562 and *φ*_3_ = 0.126 ± 0.249. The random effect for *φ*_3_ was estimated to be 0.169. While standard errors are relatively large, the purpose of the fitting process was not the parameter values per se, rather to obtain a representative model of the recovery process. The resulting model is shown in Figure S11A.

AAV following sham intervention at 5ss was modelled using the linear function

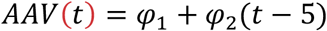

The intercept parameter *φ*_1_ was modelled with a normally-distributed random effect. Estimates for the fixed effects were *φ*_1_ = 0.356 ± 0.063 and *φ*_2_ = 0.0362 ± 0.0283; the random effect for *φ*_1_ was estimated to be 0.0378. The model is shown in Figure S11B.

In the case of sham intervention, based on linearity of Stokes flow, AAV at 3ss – 5ss was rescaled based on observed AAV at 6 ss grounded on previously-observed data on motile cilia percentage, specifically 5.57% at 3 ss, 42.9% at 4 ss, 62.1% at 5 ss and 76.3% at 6 ss (so that *AAA*(3) = (5.57/76.3) *AAA*(6) for example). In the cases of intervention at 3 ss or 4 ss, the function *AAV*_*interv*_5*ss*_(*t*) was shifted in time so that *AAV*_*interv*_3*ss*_(*t*) = *AAV*_*interv*_5*ss*_(*t* + 2), then for 3 ≤ *t* ≤ 5 rescaled based on the motile cilia percentages given above. The modelled AAV distributions for every case are given in Figure S12.

#### Signal integration and symmetry breaking

Symmetry breaking signal *S* is modelled as a weighted sum at each somite stage,

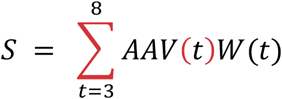

Where the parameters *W*(*t*) are the sensitivity weighting of each stage *t* = 3,…,8. The weightings are independent of the experimental situation and so are determined simultaneously across all experiment series.

A given embryo is assumed to break symmetry normally at a baseline rate of 50% in the absence of any signal at all, and at a rate of 90% if the signal exceeds a threshold (taken to be 1 in arbitrary units).

### When sampling over 1000

embryos, the abnormality rate therefore lies between 50% and 90% depending on the distribution of signal *S*. This abnormality rate is used as the success probability for the binomial model used for model fitting.

#### Signal integration model fitting

There parameters W(t) are determined through maximum likelihood estimation combined with global search, treating each experiment series with a binomial model, with success probability determined from S as described above. The log-likelihood is then maximized through global optimization, by applying the MultiStart and GlobalSearch functions (Matlab Global Optimization Toolbox). A uniform initial guess ***W***(**3**) = ***W***(**4**) = ⋯ = ***W***(**8**) = **1** is supplied to MultiStart, which executes 100000 local solver runs starting in the set [**0**, **5**]**^6^**. The output of MultiStart is then supplied to GlobalSearch as an initial guess, and the optimal value of the two solvers is taken as the best fit.

#### Restricted models

To assess whether alternative fits weighted to later somite stages could explain the data as well as the best fit to the full model, four restricted models were also assessed. These models enforced in turn *W*(3) = 0, *W*(3) = *W*(4) = 0, *W*(3) = *W*(4) = *W*(5) = 0 and *W*(3) = *W*(4) = *W*(5) = *W*(6) = 0.

## Supplemental Video S1

Close-up of fluid extraction technique employed in a 8 ss embryo.

Sequence of processed bright-field images acquired at 10x magnification, showing a micropipette extracting the liquid from the KV lumen (related to Figure 1 A-B).

## Supplemental Video S2

Close-up of fluid extraction technique employed in a 5 ss embryo.

## Supplemental Video S3

Evaluation of motile and immotile cilia distribution at 6 somite stage embryos. Injection of arl13b at 1 cell stage allows for visualization of motile versus immotile cilia. Anterior-posterior distribution was scored in 15 embryos that were later grouped as sham controls (normal situs), embryos with LR defects and without LR defects (related to Table S1).

## Supplemental Video S4

Example of a manipulated embryo that developed normal zebrafish *situs* (left heart and left liver). Sequence of processed bright-field images of a 6 ss KV is shown, oriented with anterior to the top and left to the left of the page, played at 25 fps (related to Figure 2B).

## Supplemental Video S5

Example of a manipulated embryo that developed abnormal *situs*.

Sequence of processed bright-field images of a 6 ss KV is shown, oriented with anterior to the top and left to the left of the page, played at 25 fps (related to Figure 2C).

## Supplemental Video S6

Close-up of fluid dilution technique employed in a 5 ss embryo.

Sequence of processed bright-field images acquired at 10x magnification, showing the extracted liquid from the KV lumen being mixed with a previously loaded micropipette with Danieau’s buffer (related to Figures 4A-B).

## Acknowledgments

This work was mainly supported by the Fundação para a Ciência e Tecnologia (research grant – PTDC/BEX-BID/1411/2014).

PS was funded by a PhD fellowship FCT: SFRH/BD/111611/2015.

SP was funded by a PhD fellowship FCT: SFRH/BD/130272/2017

IAT was funded by FCT Investigator IF/00082/2013, EU FP7-PEOPLE-2013-CIG (N° 818743) and by the Fundação Calouste Gulbenkian (FCG).

AG thanks to CZI Expanding Global Access to Bioimaging and to PAPIIT (IN211821) for funding support.

DJS acknowledges support from the Engineering and Physical Sciences Research Council UK via a Healthcare Technologies Challenge Award (EP/N021096/1) and a Turing Fellowship (EP/N510129/1).

SL was funded by FCT Investigator IF/00951/2012, by NOVA Medical School and by FCT CEEC-IND 2018.

Zebrafish were reared and maintained in the NMS Fish Facility, with the support from the research consortia Congento funded by LISBOA-01-0145-FEDER-022170, microscopy was performed at the IGC Advanced Imaging Facility which is supported by PPBI-POCI-01-0145-FEDER-022122; all co-financed by Lisboa Regional Operational Program (Lisboa2020), under the Portugal 2020 Partnership Agreement, through the European Regional Development Fund (ERDF) and Fundação para a Ciência e Tecnologia under the project.

The project LysoCil funded by the European Union Horizon 2020 research and innovation under grant agreement No 811087 supported SL, PS and SP travelling to the FASEB cilia meeting in the USA in 2019, where this work was presented and discussed with peers. We thank Sérgio Dias for reviewing the final version of the manuscript.

## Author contributions

Conceptualization: SSL, IT, PS, DS

Methodology: PS, SP, IT, DS

Investigation: SSL, PS, DS

Visualization: SSL, DS,

Funding acquisition: SSL

Project administration: SSL

Supervision: SSL, IT, AG, DS

Writing – original draft: PS, SSL

Writing – review & editing: SSL, IT, SP, PS, DS, AG

## Data and materials availability

All data are available in the main text or the supplementary materials.

## References

Agarwal, K. et al. (2015) ‘Analysis of Exosome Release as a Cellular Response to MAPK Pathway Inhibition’, Langmuir, 31(19), pp. 5440–5448. doi:10.1021/acs.langmuir.5b00095.

Asadnia, M. et al. (2016) ‘From Biological Cilia to Artificial Flow Sensors: Biomimetic Soft Polymer Nanosensors with High Sensing Performance’, Scientific Reports, 6(1), p. 32955. doi:10.1038/srep32955.

Bataille, S. et al. (2011) ‘Association of PKD2 (polycystin 2) mutations with left-right laterality defects’, American Journal of Kidney Diseases, 58(3), pp. 456–460. doi:10.1053/j.ajkd.2011.05.015.

Brown, K.T. and Flaming, D.G. (1974) ‘Beveling of fine micropipette electrodes by a rapid precision method’, Science [Preprint]. doi:10.1126/science.185.4152.693.

Cartwright, J.H.E., Piro, O. and Tuval, I. (2020) ‘Chemosensing versus mechanosensing in nodal and Kupffer’s vesicle cilia and in other left–right organizer organs’, Philosophical Transactions of the Royal Society B: Biological Sciences, 375(1792), p. 20190566. doi:10.1098/rstb.2019.0566.

Chiu, Y.-J. et al. (2016) ‘A Single-Cell Assay for Time Lapse Studies of Exosome Secretion and Cell Behaviors’, Small, 12(27), pp. 3658–3666. doi:10.1002/smll.201600725.

Chung, W.-S. and Stainier, D.Y.R. (2008) ‘Intra-Endodermal Interactions Are Required for Pancreatic β Cell Induction’, Developmental Cell, 14(4), pp. 582–593. doi:10.1016/j.devcel.2008.02.012.

Delling, M. et al. (2016) ‘Primary cilia are not calcium-responsive mechanosensors’, Nature, 531(7596), pp. 656–660. doi:10.1038/nature17426.

Essner, J.J. et al. (2005) ‘Kupffer’s vesicle is a ciliated organ of asymmetry in the zebrafish embryo that initiates left-right development of the brain, heart and gut.’, Development (Cambridge, England), 132(6), pp. 1247–1260. doi:10.1242/dev.01663.

Ferreira, R.R. et al. (2017) ‘Physical limits of flow sensing in the left-right organizer’, eLife, 6. doi:10.7554/eLife.25078.

Le Fevre, A. et al. (2020) ‘Compound heterozygous Pkd1l1 variants in a family with two fetuses affected by heterotaxy and complex Chd’, European Journal of Medical Genetics, 63(2), p. 103657. doi:10.1016/j.ejmg.2019.04.014.

Field, S. et al. (2011) ‘Pkd1l1 establishes left-right asymmetry and physically interacts with Pkd2’, Development, 138(6), pp. 1131–1142. doi:10.1242/dev.058149.

Gallagher, M.T. et al. (2019) ‘Rapid sperm capture: high-throughput flagellar waveform analysis’, Human Reproduction [Preprint]. doi:10.1093/humrep/dez056.

Gokey, J.J. et al. (2016) ‘Kupffer’s vesicle size threshold for robust left-right patterning of the zebrafish embryo’, Developmental Dynamics, 245(1), pp. 22–33. doi:10.1002/dvdy.24355.

Hashimoto, H. et al. (2004) ‘The Cerberus/Dan-family protein Charon is a negative regulator of Nodal signaling during left-right patterning in zebrafish.’, Development (Cambridge, England), 131(8), pp. 1741–1753. doi:10.1242/dev.01070.

Hojo, M. et al. (2007) ‘Right-elevated expression of charon is regulated by fluid flow in medaka Kupffer’s vesicle’, Development Growth and Differentiation, 49(5), pp. 395–405. doi:10.1111/j.1440-169X.2007.00937.x.

Jacinto, R. et al. (2021) ‘Pkd2 Affects Cilia Length and Impacts LR Flow Dynamics and Dand5’, Frontiers in Cell and Developmental Biology, 9. doi:10.3389/fcell.2021.624531.

Juan, T. et al. (2018) ‘Myosin1D is an evolutionarily conserved regulator of animal left–right asymmetry’, Nature Communications, 9(1), p. 1942. doi:10.1038/s41467-018-04284-8.

Kamura, K. et al. (2011) ‘Pkd1l1 complexes with Pkd2 on motile cilia and functions to establish the left-right axis.’, Development (Cambridge, England), 138(6), pp. 1121–1129. doi:10.1242/dev.058271.

Kikuchi, Y. et al. (2004) ‘Notch Signaling Can Regulate Endoderm Formation in Zebrafish’, Developmental Dynamics, 229(4), pp. 756–762. doi:10.1002/dvdy.10483.

Kimmel, C.B. et al. (1995) ‘Stages of embryonic development of the zebrafish.’, Developmental dynamics : an official publication of the American Association of Anatomists, 203(3), pp. 253–310. doi:10.1002/aja.1002030302.

Lopes, S.S. et al. (2010) ‘Notch signalling regulates left-right asymmetry through ciliary length control.’, Development (Cambridge, England), 137(21), pp. 3625–3632. doi:10.1242/dev.054452.

Maerker, M. et al. (2021) ‘Bicc1 and Dicer regulate left-right patterning through post-transcriptional control of the Nodal inhibitor Dand5’, Nature Communications, 12(1), p. 5482. doi:10.1038/s41467-021-25464-z.

Marques, S. et al. (2004) ‘The activity of the Nodal antagonist Cerl-2 in the mouse node is required for correct L/R body axis’, Genes and Development, 18(19), pp. 2342–2347. doi:10.1101/gad.306504.

McGrath, J. et al. (2003) ‘Two populations of node monocilia initiate left-right asymmetry in the mouse’, Cell, 114(1), pp. 61–73. doi:10.1016/S0092-8674(03)00511-7.

Minegishi, K. et al. (2021) ‘Fluid flow-induced left-right asymmetric decay of Dand5 mRNA in the mouse embryo requires a Bicc1-Ccr4 RNA degradation complex’, Nature Communications, 12(1), p. 4071. doi:10.1038/s41467-021-24295-2.

Mizuno, K. et al. (2020) ‘Role of Ca ^2+^ transients at the node of the mouse embryo in breaking of left-right symmetry’, Science Advances, 6(30). doi:10.1126/sciadv.aba1195.

Navis, A., Marjoram, L. and Bagnat, M. (2013) ‘Cftr controls lumen expansion and function of Kupffer’s vesicle in zebrafish’, 1712, pp. 1703–1712. doi:10.1242/dev.091819.

Nonaka, S. et al. (1998) ‘Randomization of left-right asymmetry due to loss of nodal cilia generating leftward flow of extraembryonic fluid in mice lacking KIF3B motor protein’, Cell, 95(6), pp. 829–837. doi:10.1016/S0092-8674(00)81705-5.

R. Ferreira, R. et al. (2019) ‘The cilium as a force sensor-myth versus reality’, Journal of Cell Science, 132(14). doi:10.1242/jcs.213496.

Roxo-Rosa, M. et al. (2015) ‘The zebrafish Kupffer’s vesicle as a model system for the molecular mechanisms by which the lack of Polycystin-2 leads to stimulation of CFTR’, Biol Open, 4(11), pp. 1356–1366. doi:10.1242/bio.014076.

Roxo-Rosa, M. and Santos Lopes, S. (2020) ‘The Zebrafish Kupffer’s Vesicle: A Special Organ in a Model Organism to Study Human Diseases’, in Zebrafish in Biomedical Research. IntechOpen. doi:10.5772/intechopen.88266.

Sampaio, P. et al. (2014) ‘Left-right organizer flow dynamics: How much cilia activity reliably yields laterality?’, Developmental Cell, 29(6), pp. 716–728. doi:10.1016/j.devcel.2014.04.030.

Schweickert, A. et al. (2007) ‘Cilia-Driven Leftward Flow Determines Laterality in Xenopus’, Current Biology, 17(1), pp. 60–66. doi:10.1016/j.cub.2006.10.067.

Schweickert, A. et al. (2010) ‘The Nodal Inhibitor Coco Is a Critical Target of Leftward Flow in Xenopus’, Current Biology, 20(8), pp. 738–743. doi:10.1016/j.cub.2010.02.061.

Shinohara, K. et al. (2012) ‘Two rotating cilia in the node cavity are sufficient to break left–right symmetry in the mouse embryo’, Nature Communications, 3, p. 622. doi:10.1038/ncomms1624.

Shiratori, H. and Hamada, H. (2006) ‘The left-right axis in the mouse: from origin to morphology.’, Development (Cambridge, England), 133(11), pp. 2095–104. doi:10.1242/dev.02384.

Smith, D.J., Montenegro-Johnson, T.D. and Lopes, S.S. (2014) ‘Organized chaos in Kupffer’s vesicle: how a heterogeneous structure achieves consistent left-right patterning’, Bioarchitecture, 4(3), pp. 119–125. doi:10.4161/19490992.2014.956593.

Supp, D.M. et al. (1999) ‘Targeted deletion of the ATP binding domain of left-right dynein confirms its role in specifying development of left-right asymmetries.’, Development (Cambridge, England), 126(23), pp. 5495–504. Available at: http://www.pubmedcentral.nih.gov/articlerender.fcgi?artid=1797880&tool=pmcentrez&rendertype=abstract.

Takao, D. et al. (2013) ‘Asymmetric distribution of dynamic calcium signals in the node of mouse embryo during left–right axis formation’, Developmental Biology, 376(1), pp. 23–30. doi:10.1016/j.ydbio.2013.01.018.

Tanaka, Y., Okada, Y. and Hirokawa, N. (2005) ‘FGF-induced vesicular release of Sonic hedgehog and retinoic acid in leftward nodal flow is critical for left-right determination.’, Nature, 435(7039), pp. 172–177. doi:10.1038/nature03494.

Tavares, B. et al. (2017) ‘Notch/Her12 signalling modulates, motile/immotile cilia ratio downstream of Foxj1a in zebrafish left-right organizer’, eLife, 6, p. e25165. doi:10.7554/eLife.25165.

Wang, G. et al. (2011) ‘The Rho kinase Rock2b establishes anteroposterior asymmetry of the ciliated Kupffer’s vesicle in zebrafish.’, Development (Cambridge, England), 138(1), pp. 45–54. doi:10.1242/dev.052985.

Wilson, K.S. et al. (2015) ‘Dynein-deficient flagella respond to increased viscosity with contrasting changes in power and recovery strokes’, Cytoskeleton, 72(9), pp. 477–490. doi:10.1002/cm.21252.

Yuan, S. et al. (2015) ‘Intraciliary calcium oscillations initiate vertebrate left-right asymmetry’, Current Biology, 25(5), pp. 556–567. doi:10.1016/j.cub.2014.12.051.

