## supplemental figures for "Fluid extraction from the left-right organizer uncovers mechanical properties needed for symmetry breaking"

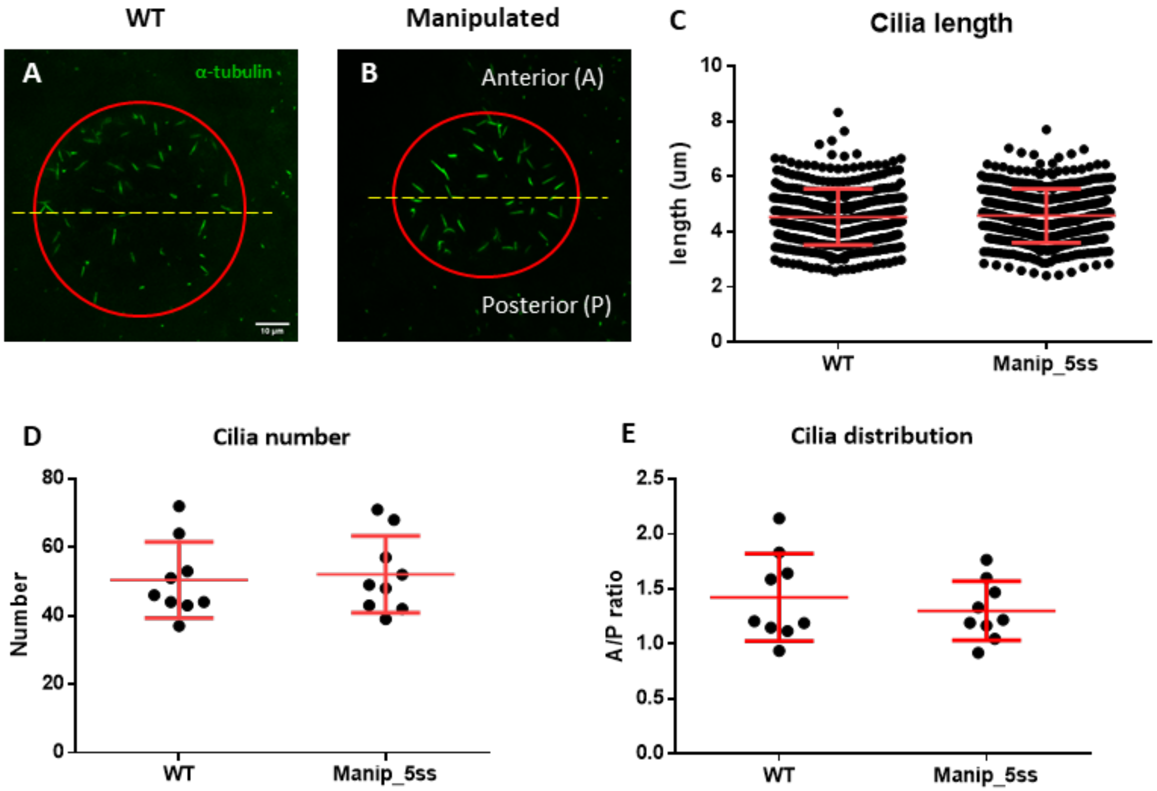


**Figure S1 – Manipulated embryos develop normal LRO cilia evaluated by immunofluorescence.** Cilia number, length and anterior-posterior ratio of cilia distribution in WT and fluid extracted embryos – (A-B): Acetylated α-tubulin (in green) immunostaining examples showing LRO labelled cilia on WT and manipulated embryos. A number of 388 cilia in 8 WT embryos and 399 cilia in 8 manipulated embryos were measured in 3D (C). Mean cilia number (D) and A/P ratio (E) are displayed per embryo. Student's t-test was used to assess differences between groups.

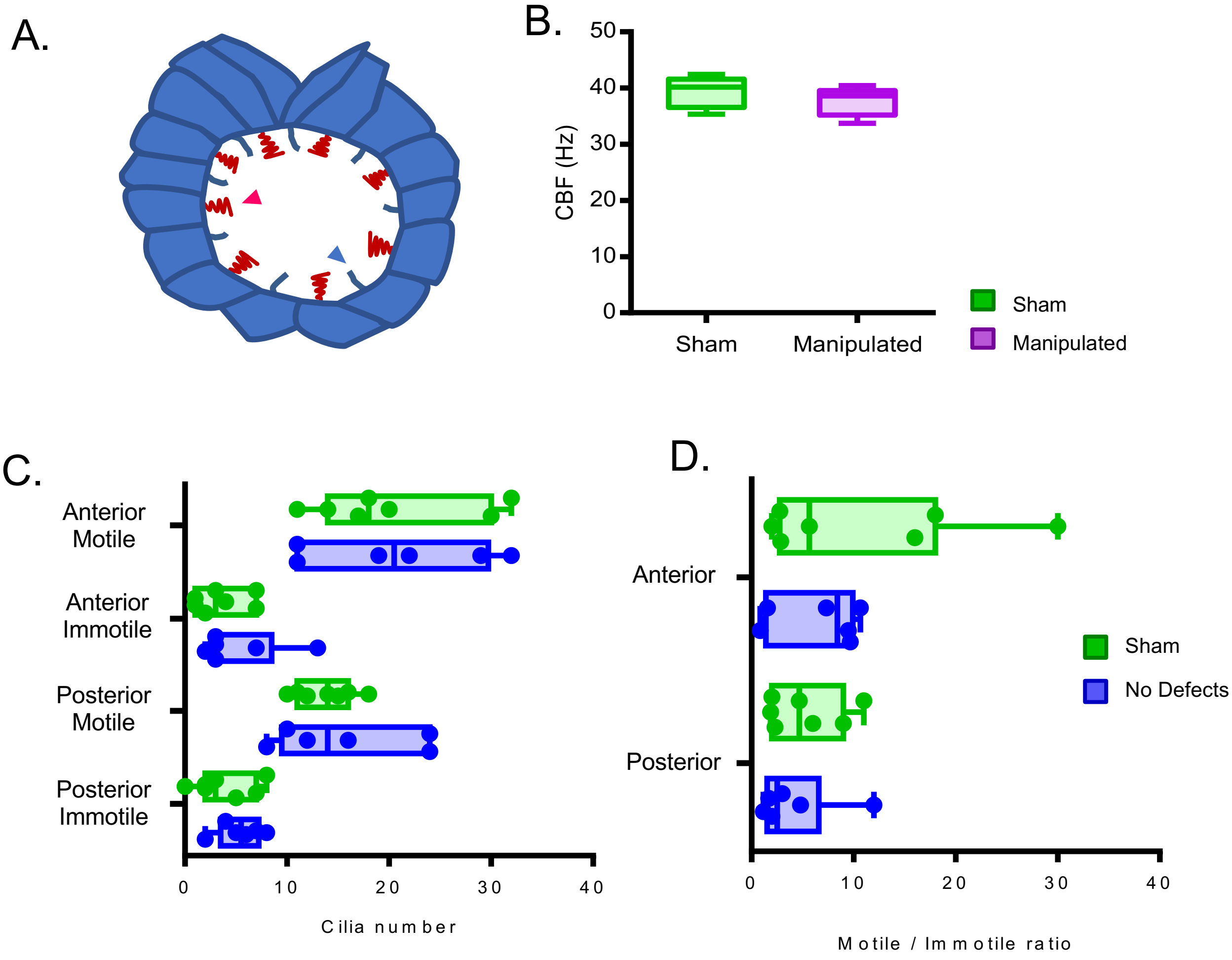


**Figure S2 – Cilia Beat Frequency and motile/ immotile cilia ratio do not change in manipulated embryos evaluated by live imaging.** (A) Diagram highlighting motile and immotile cilia intercalated in the LRO cells. (B) CBF of the motile beating cilia that could be visualized by bright field live microscopy in the Sham control embryos versus manipulated embryos (C) Total number of motile and immotile cilia located anteriorly and posteriorly between sham controls and embryos without LR defects. (D) Motile to immotile cilia ratio in the anterior and posterior LRO of Sham control embryos versus embryos that later developed without LR defects. Fisher's exact test was used to assess differences between treatments using pooled data from 6 embryos per treatment. Embryos were manipulated for LRO fluid extracted at 5 ss and were imaged at 6 ss. CBF: cilia beat frequency.

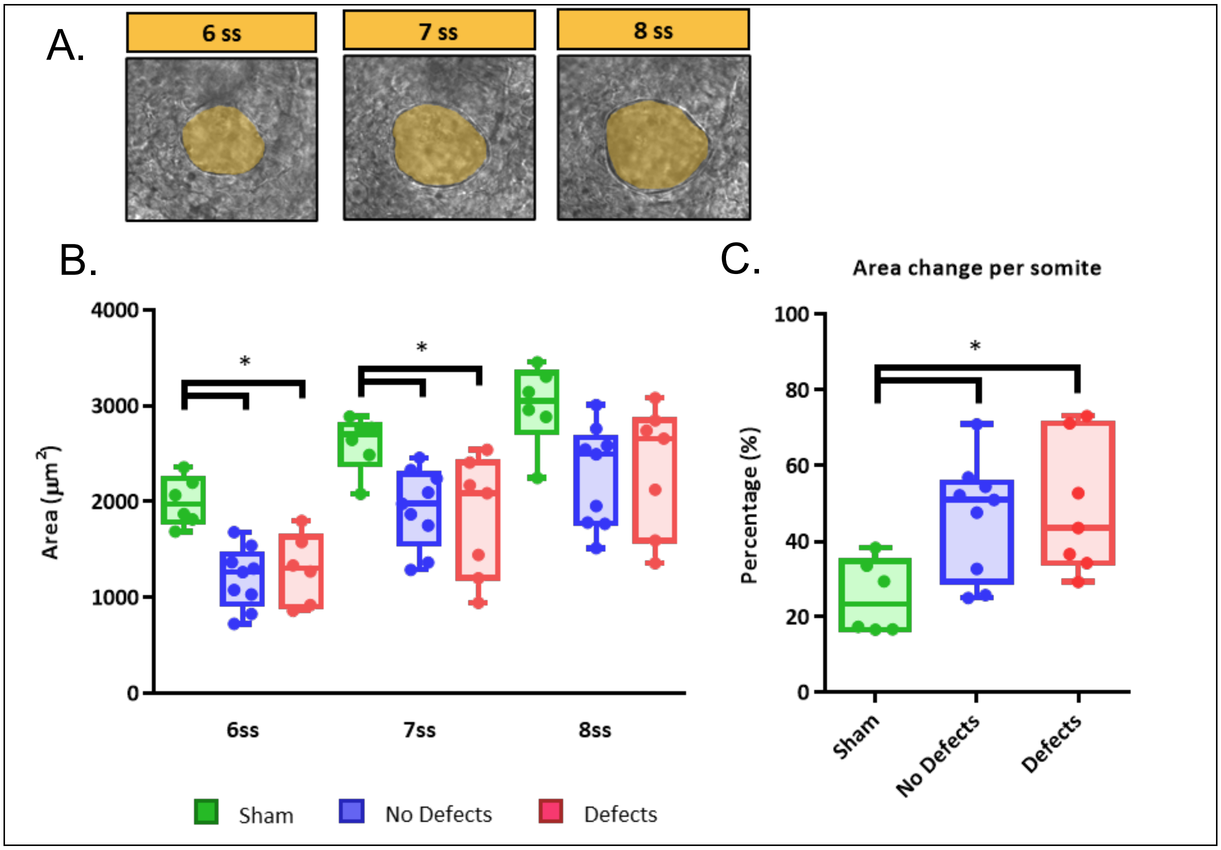


**Figure S3 – LRO areas from recovering embryos that develop left-right defects are not different from embryos that have a normal development.** (A) LRO areas at 6-8 ss following extraction at 5 ss (B) Quantifications of LRO area of the three different groups (“Sham” control, “No Defects” and “Defects” group) in manipulated embryos during LRO lumen area recovery from 6 ss to 8 ss. (C) Area change per somite. Mann-Whitney U test p-value <0.05.

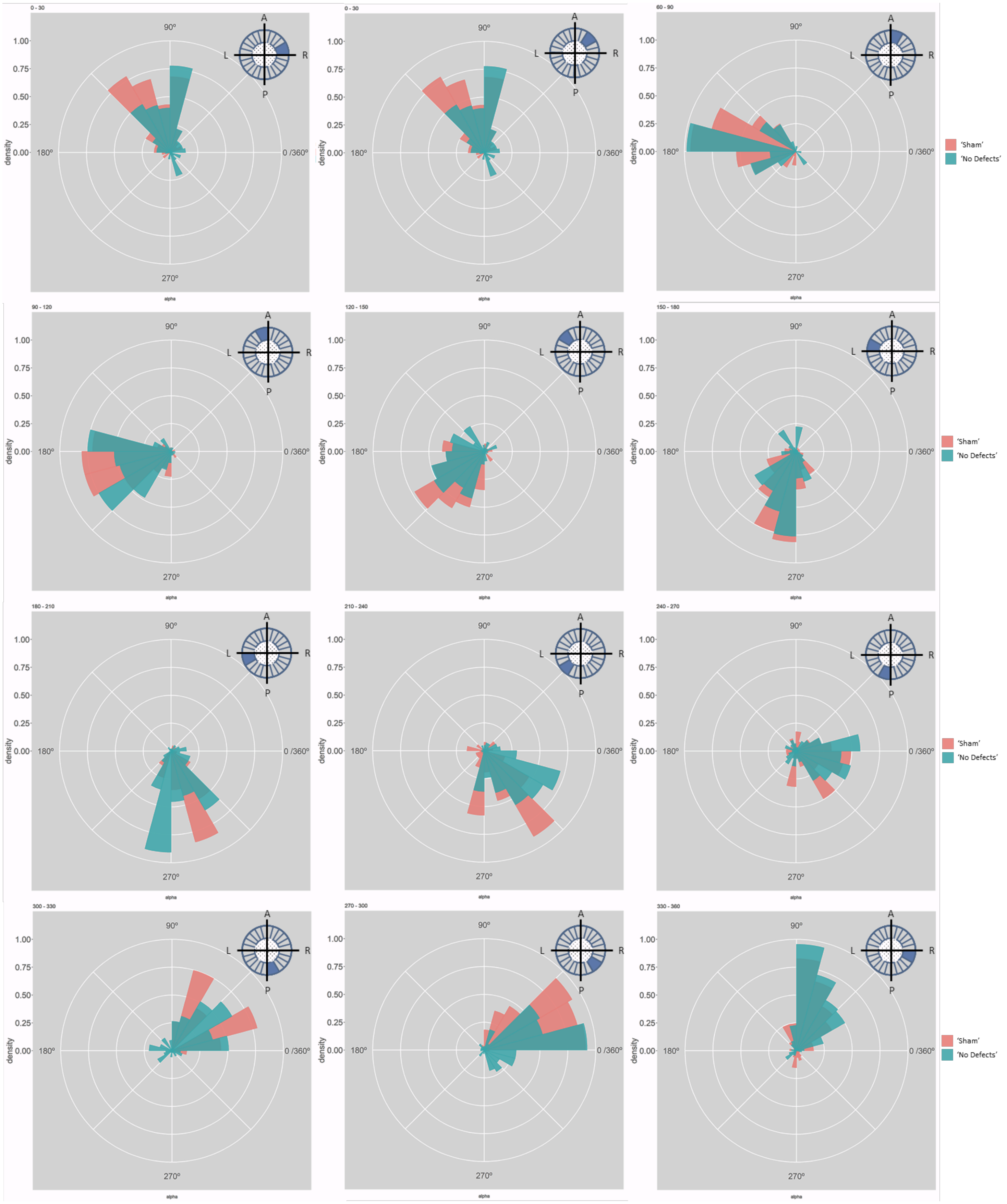
0 - 30
0 - 30
60 - 90
90 - 120
120 - 150
150 - 180
180 - 210
210 - 240
240 - 270
300 - 330
270 - 300
330 - 360
density
alpha
1.00
0.75
0.50
0.25
0.00
90º
180º
270º
0 /360º
A
P
L
R
'Sham'
'No Defects'

**Figure S4– Particle movement direction at different regions of the LRO for 'Sham' and 'No Defects' groups at 6 ss.** Each plot represents the pooled tracked trajectory of a moving particle at even given point in time. Trajectories were plotted from the outermost area of the LRO (radius > 0.5 from a maximum of 1). Kolmogorov-Smirnov test was used from comparing trajectory distribution between the two groups. ss: somite stage. LRO: left-right organizer.

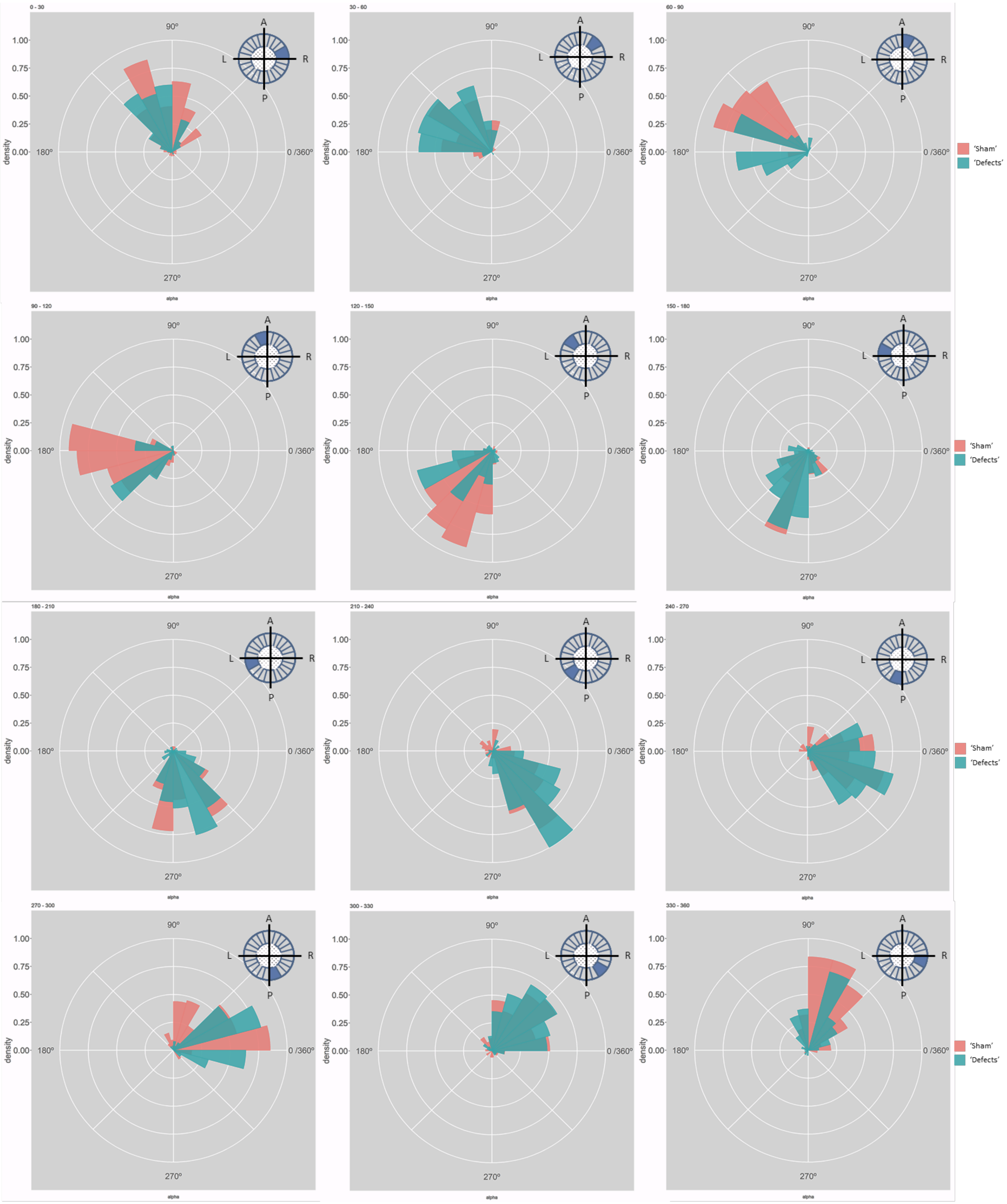

0 - 30
30 - 60
60 - 90
90 - 120
120 - 150
150 - 180
180 - 210
210 - 240
240 - 270
270 - 300
300 - 330
330 - 360
density
1.00
0.75
0.50
0.25
0.00
90º
180º
270º
0 /360º
alpha
A
P
L
R
'Sham'
'Defects'

**Figure S5** – **Particle movement direction at different regions of the LRO for 'Sham' and 'No Defects' groups at 7 ss.** Each plot represents the pooled tracked trajectory of a moving particle at even given point in time. Trajectories were plotted from the outermost area of the KV's (radius > 0.5 from a maximum of 1). Kolmogrov-Smirnov test was used from comparing trajectory distribution between the two groups. ss: somite stage; LRO: left-right organizer.

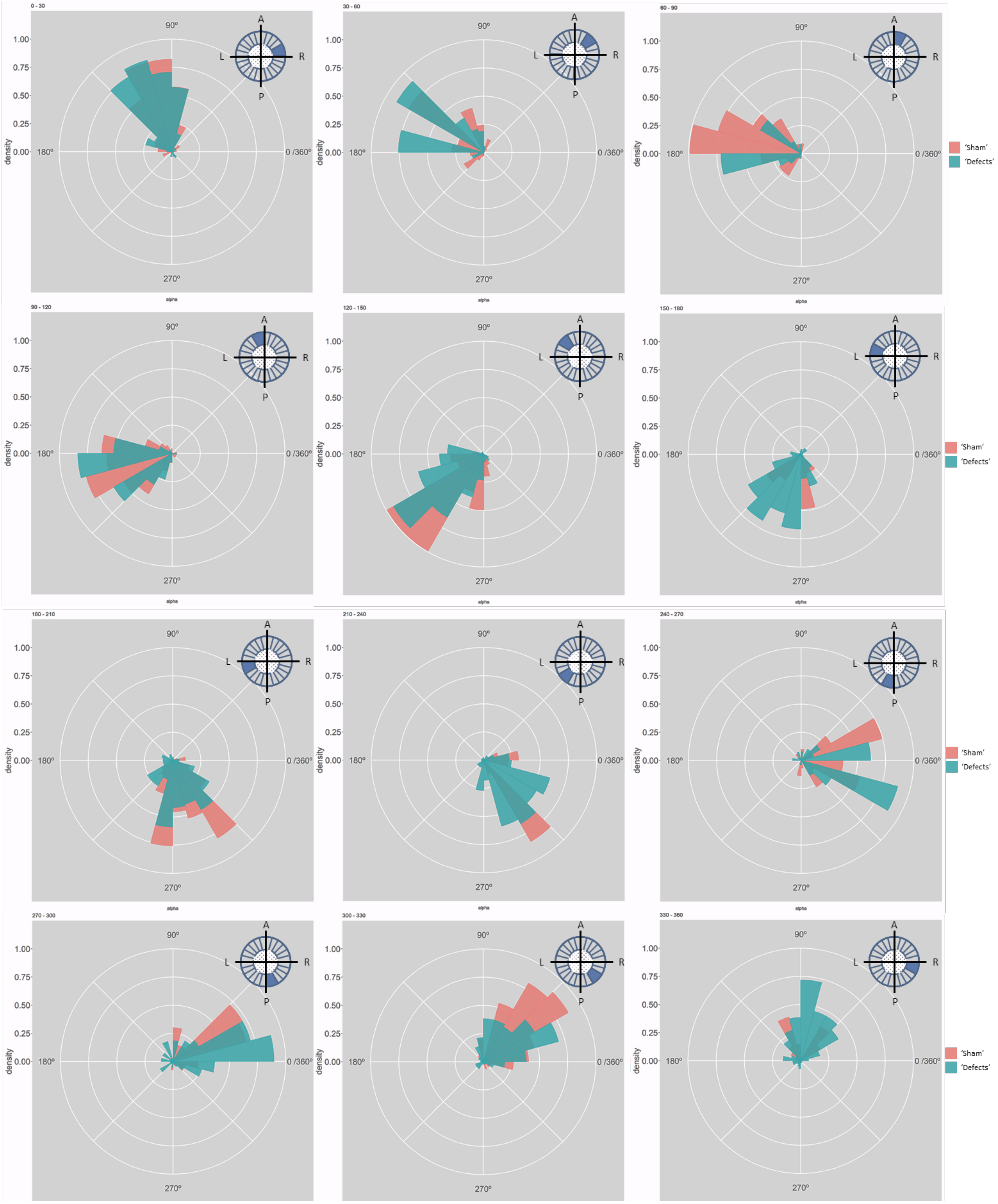
0 - 30
30 - 60
60 - 90
90 - 120
120 - 150
150 - 180
180 - 210
210 - 240
240 - 270
270 - 300
300 - 330
330 - 360
1.00
0.75
0.50
0.25
0.00
density
alpha
90º
180º
270º
0 /360º
A
L
R
P
'Sham'
'Defects'

**Figure S6– Particle movement direction at different regions of the LRO for 'Sham' and 'No Defects' groups at 8ss**. Each plot represents the pooled tracked trajectory of a moving particle at even given point in time. Trajectories were plotted from the outermost area of the KV's (radius > 0.5 from a maximum of 1). Kolmogorov-Smirnov test was used from comparing trajectory distribution between the two groups. ss: somite stage. LRO: left-right organizer.

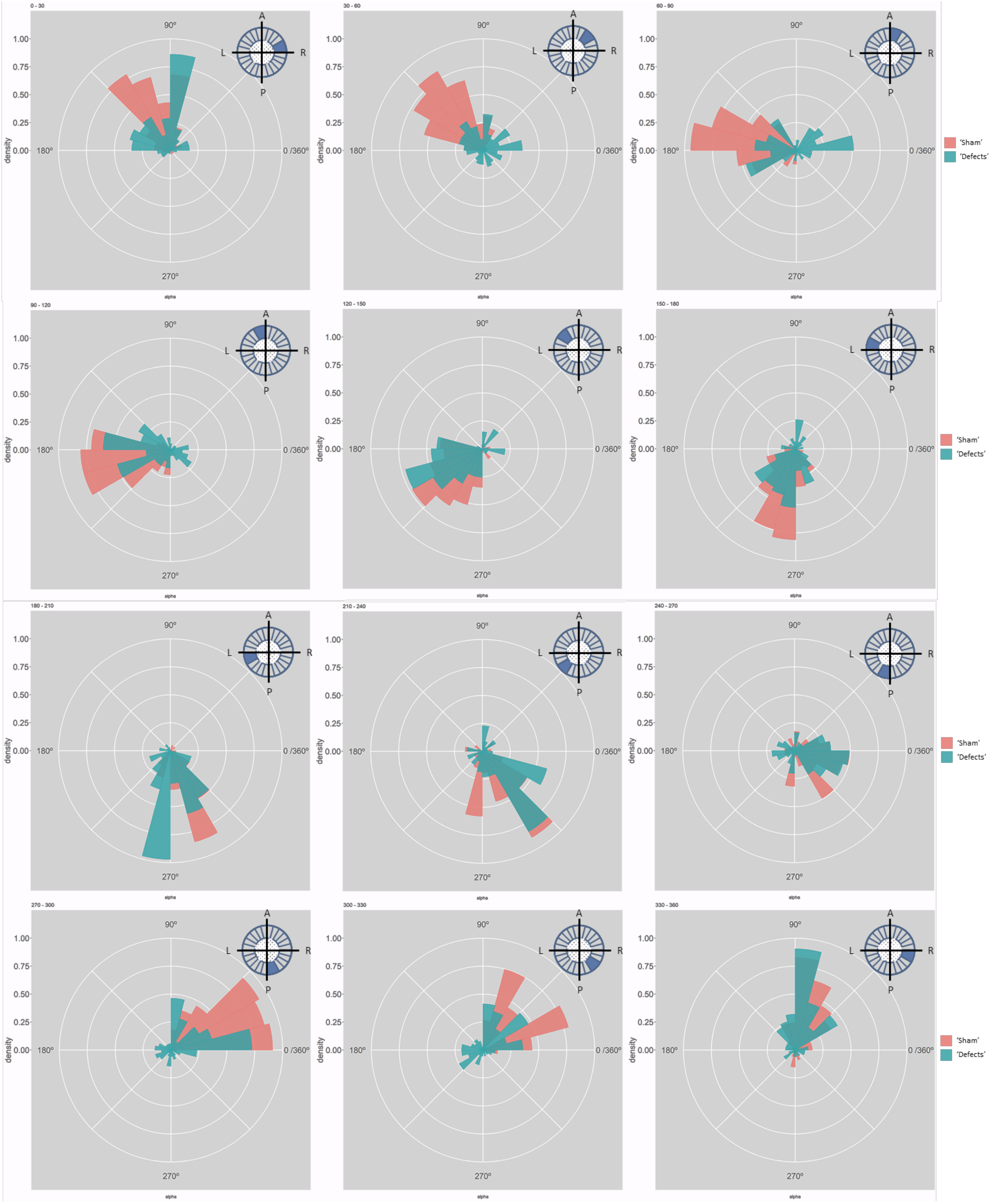
0 - 30
30 - 60
60 - 90
90 - 120
120 - 150
150 - 180
180 - 210
210 - 240
240 - 270
270 - 300
300 - 330
330 - 360
90º
180º
0 /360º
270º
density
alpha
A
P
L
R
'Sham'
'Defects'

**Figure S7 – Particle movement direction at different regions of the KV for 'Sham' and 'Defects' groups at 6 ss.** Each plot represents the pooled tracked trajectory of a moving particle at even given point in time. Trajectories were plotted from the outermost area of the KV's (radius > 0.5 from a maximum of 1) Kolmogorov-Smirnov test was used from comparing trajectory distribution between the two groups. ss: somite stage. LRO: left-right organizer.

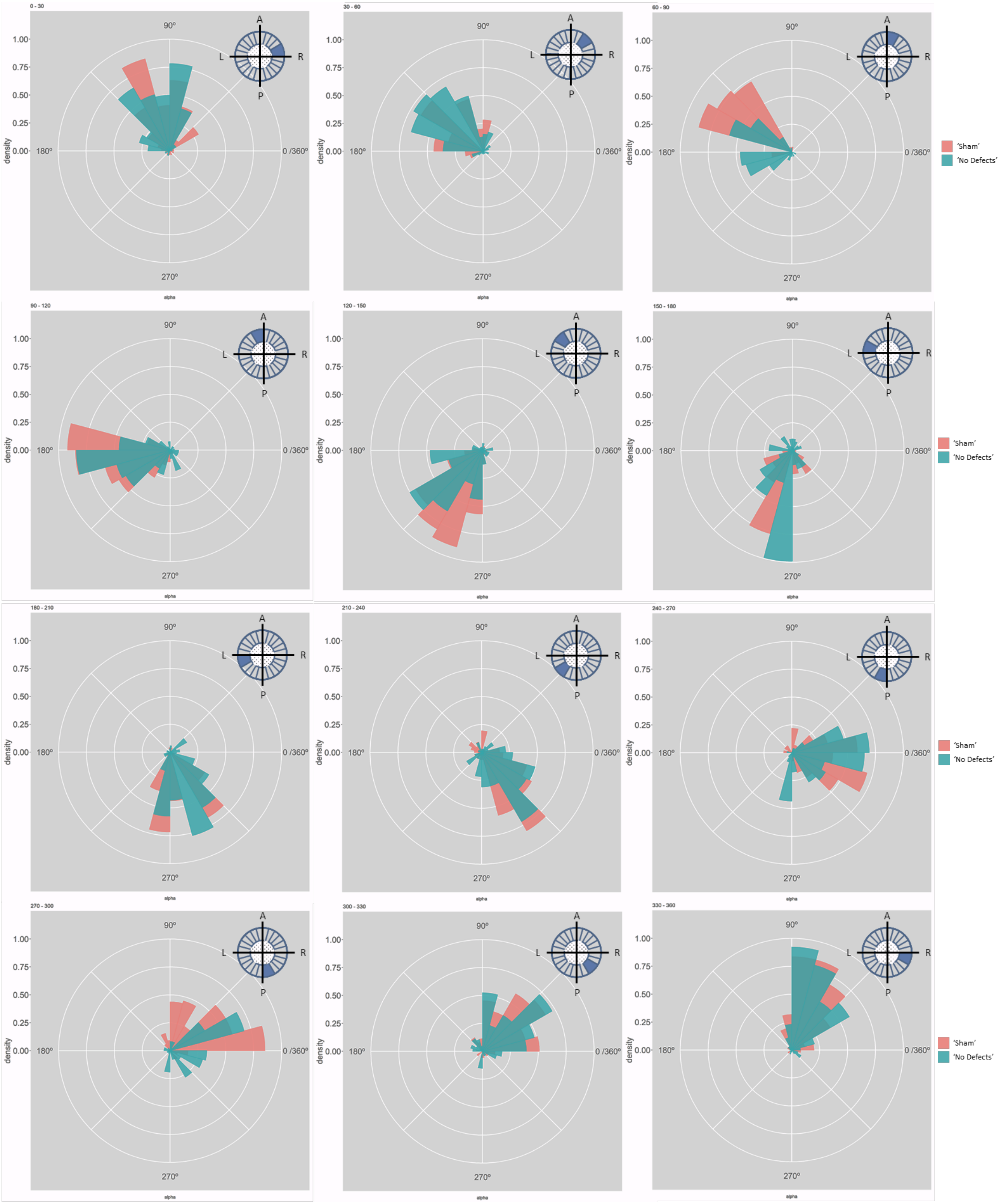
0 - 30
30 - 60
60 - 90
90 - 120
120 - 150
150 - 180
180 - 210
210 - 240
240 - 270
270 - 300
300 - 330
330 - 360
90°
180°
270°
0 /360°
1.00
0.75
0.50
0.25
0.00
density
alpha
A
L
R
P
'Sham'
'No Defects'

**Figure S8– Particle movement direction at different regions of the LRO for 'Sham' and 'Defects' groups at 7 ss.** Each plot represents the pooled tracked trajectory of a moving particle at even given point in time. Trajectories were plotted from the outermost area of the KV's (radius > 0.5 from a maximum of 1). Kolmogorov-Smirnov test was used from comparing trajectory distribution between the two groups. ss: somite stage. LRO: left-right organizer.

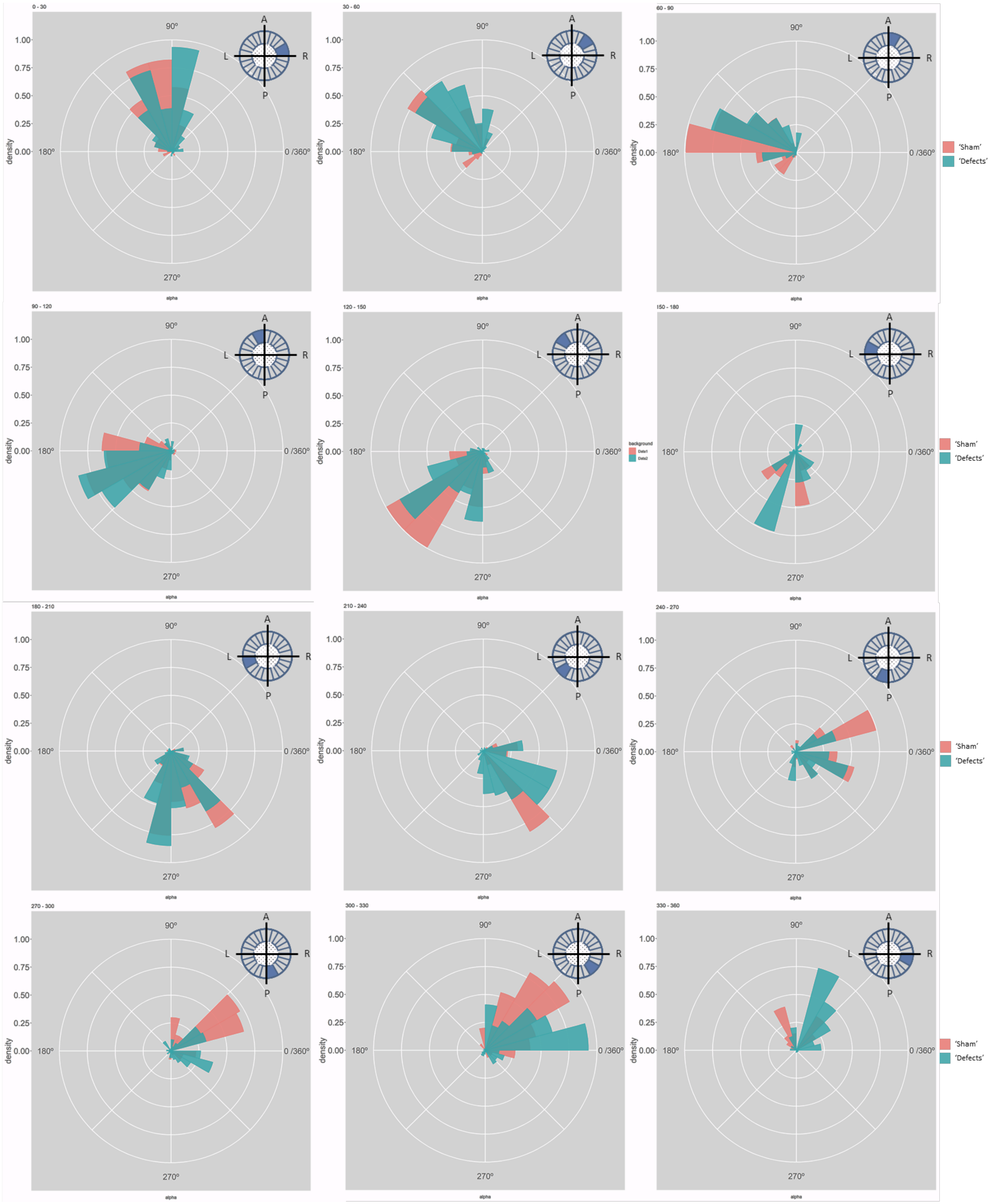
0 - 30
30 - 60
60 - 90
90 - 120
120 - 150
150 - 180
180 - 210
210 - 240
240 - 270
270 - 300
300 - 330
330 - 360
density
alpha
1.00
0.75
0.50
0.25
0.00
90º
180º
0 /360º
270º
A
P
L
R
'Sham'
'Defects'
background

**Figure S9**– **Particle movement direction at different regions of the LRO for ‘Sham’ and ‘Defects’ groups at 8 ss.** Each plot represents the pooled tracked trajectory of a moving particle at even given point in time. Trajectories were plotted from the outermost area of the KV’s (radius > 0.5 from a maximum of 1). Kolmogorov-Smirnov test was used from comparing trajectory distribution between the two groups. ss: somite stage. LRO: left-right organizer.

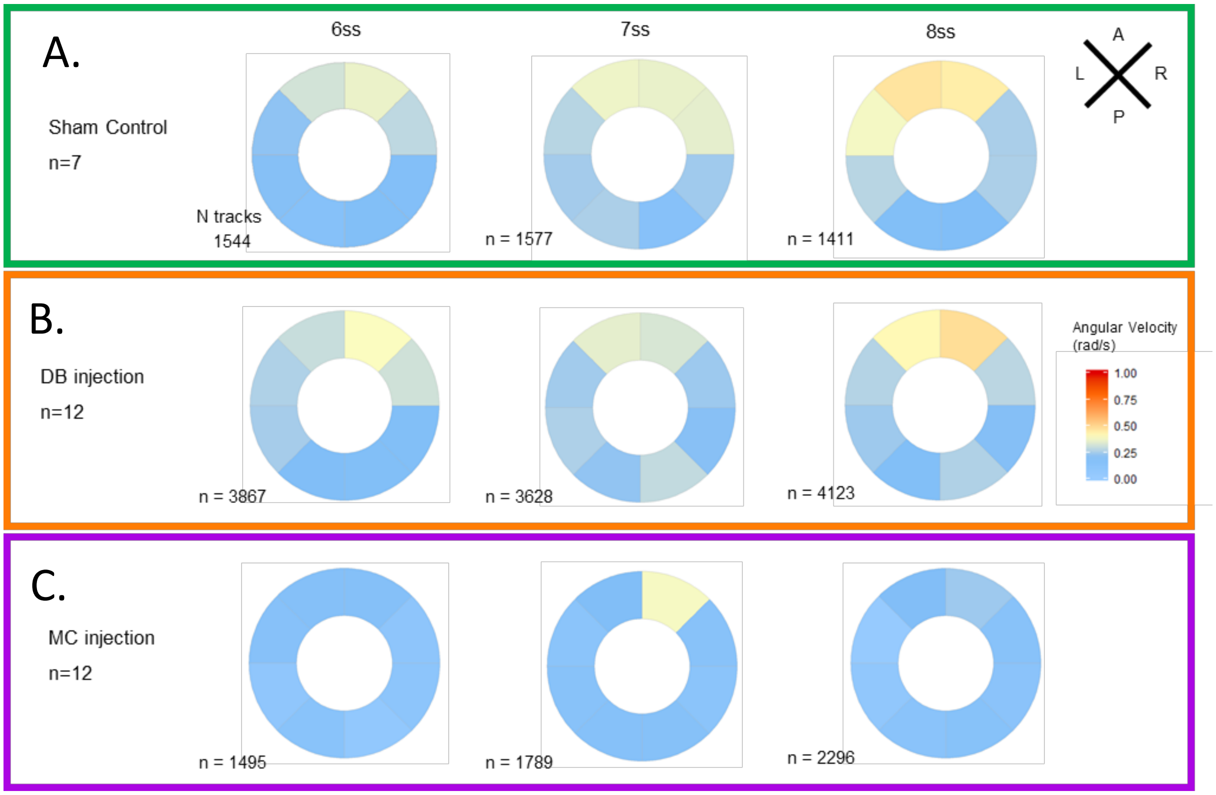


**Figure S10 –– LRO flow dynamics is only affected when fluid dilution alters fluid viscosity** (A) Angular velocity polar plots for 6 ss, 7 ss and 8 ss for the three different groups (A) “Sham” control, (B) “DB dilution” group and (C) “MC dilution” group. Number of tracks refers to the number of particle trajectories identified for the quantifications and respective angular velocity plots. Colour code on polar plots refers to the median angular velocity for all pooled embryos. LRO: left-right organizer. DB: Danieau buffer. MC: methylcellulose.

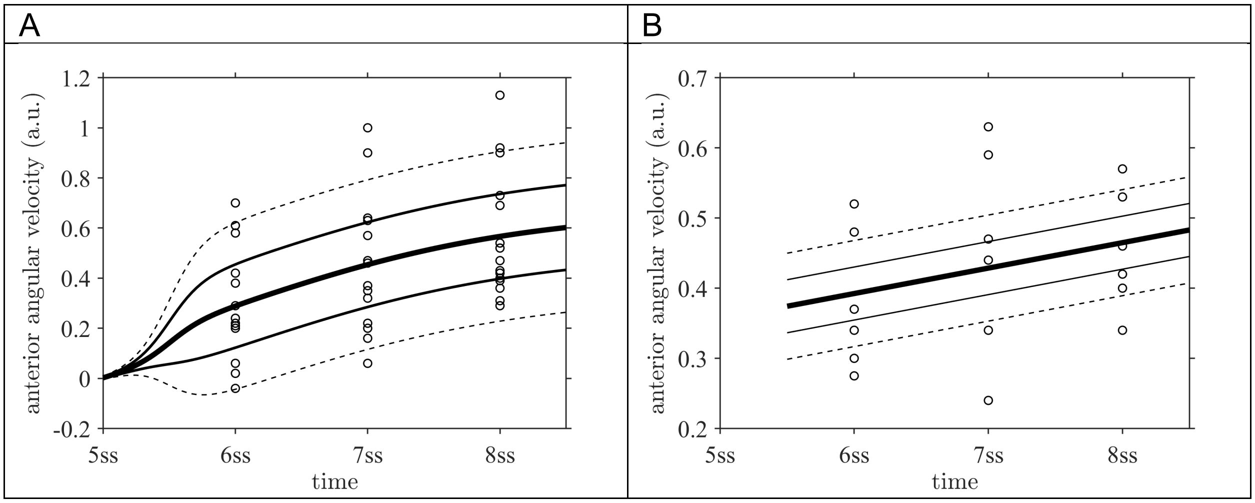


**Figure S11– Mixed effects model fits for the distributions of anterior angular velocity** to (A) data for sham intervention, (B) data for suction intervention at 5 ss. Central line is fixed effect, outer pairs of lines are +/- 1 and +/- 2 standard deviations in normally distribution random effects. ss : somite stage.

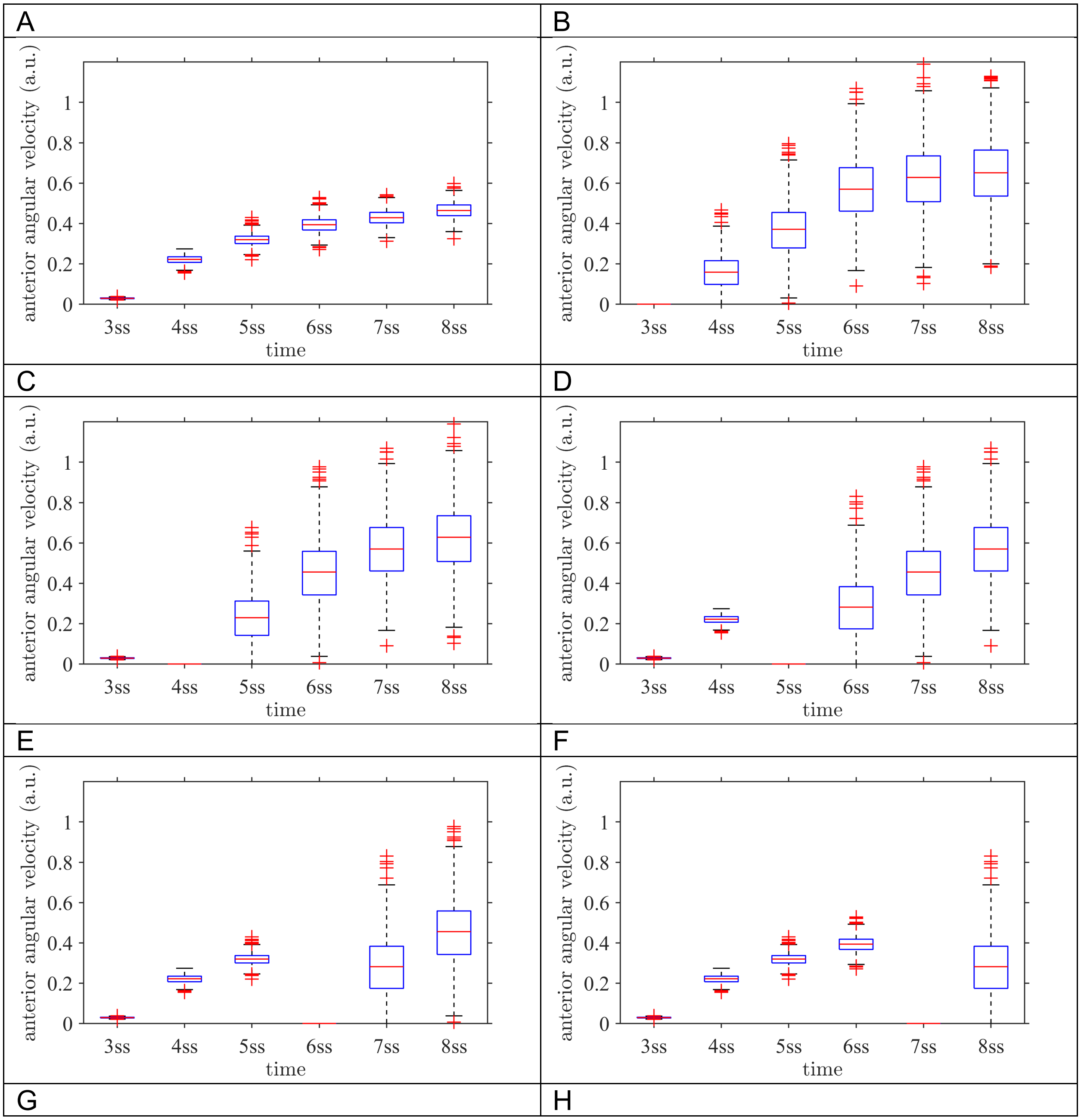
A
anterior angular velocity (a.u.)
0
0.2
0.4
0.6
0.8
1
3ss
4ss
5ss
6ss
7ss
8ss
time
B
anterior angular velocity (a.u.)
0
0.2
0.4
0.6
0.8
1
3ss
4ss
5ss
6ss
7ss
8ss
time
C
anterior angular velocity (a.u.)
0
0.2
0.4
0.6
0.8
1
3ss
4ss
5ss
6ss
7ss
8ss
time
D
anterior angular velocity (a.u.)
0
0.2
0.4
0.6
0.8
1
3ss
4ss
5ss
6ss
7ss
8ss
time
E
anterior angular velocity (a.u.)
0
0.2
0.4
0.6
0.8
1
3ss
4ss
5ss
6ss
7ss
8ss
time
F
anterior angular velocity (a.u.)
0
0.2
0.4
0.6
0.8
1
3ss
4ss
5ss
6ss
7ss
8ss
time
G
H

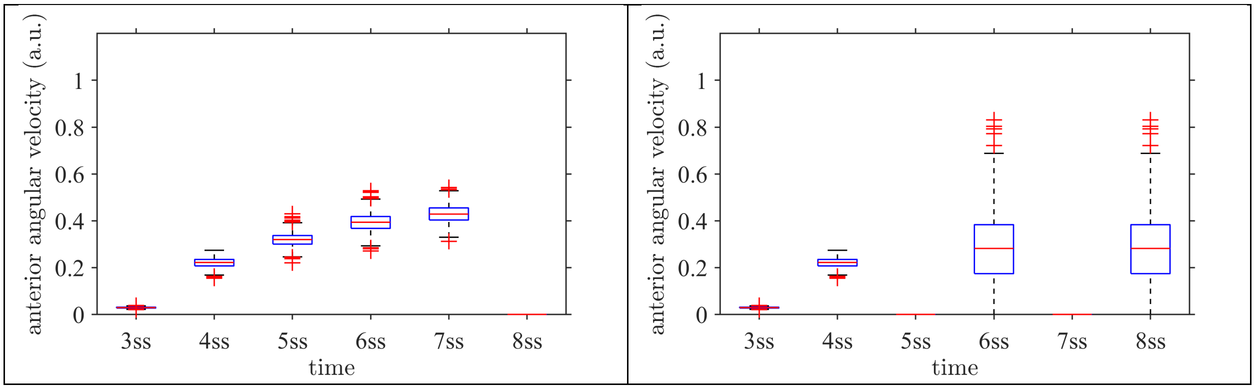


**Figure S12 - Anterior angular velocity distributions for all modelled experimental interventions**. (A) sham, (B) 3 ss intervention, (C) 4 ss intervention, (D) 5 ss intervention, (E) 6 ss intervention, (F) 7 ss intervention, (G) 8 ss intervention, (H) both 5 ss and 7 ss intervention. These distributions are combined with stage weightings to fit the normal/defect percentages for each experiment series. ss: somite stage.

### SUPPLEMENTAL TABLE:

**Table S1. Motile and immotile cilia distribution at 5 somite stage embryos**. Analyses of cilia motility per LRO anterior and posterior halves.

| | Embryos | Anterior | | Posterior | | Total motile cilia | Total cilia |
|---|---|---|---|---|---|---|---|
| | | Motile | Immotile | Motile | Immotile | | |
| Sham | E1 | 32 | 2 | 18 | 2 | 50 | 54 |
| | E2 | 20 | 7 | 15 | 8 | 35 | 50 |
| | E3 | 11 | 4 | 10 | 5 | 21 | 30 |
| | E4 | 18 | 1 | 11 | 0 | 29 | 30 |
| | E5 | 17 | 3 | 12 | 2 | 29 | 34 |
| | E6 | 30 | 1 | 16 | 7 | 46 | 54 |
| | E7 | 14 | 7 | 14 | 3 | 28 | 38 |
| No LR Defects | E1 | 22 | 3 | 16 | 8 | 38 | 49 |
| | E2 | 11 | 7 | 10 | 6 | 21 | 34 |
| | E3 | 19 | 2 | 12 | 4 | 31 | 37 |
| | E4 | 21 | 2 | 24 | 5 | 45 | 52 |
| | E5 | 32 | 3 | 24 | 2 | 56 | 61 |
| | E6 | 11 | 13 | 8 | 7 | 19 | 39 |
| LR Defects | E1 | 3 | 19 | 3 | 22 | 6 | 47 |
| | E2 | 15 | 10 | 12 | 3 | 27 | 40 |
